# Distinct druggable biological processes in early-onset prostate cancer

**DOI:** 10.1101/2025.04.25.650707

**Authors:** Stephanie T Schmidt, Zsofia Kote-Jarai, Adrian Larkeryd, Burga Ozer, Questa Karlsson, James Campbell, Sue Merson, Nenning Dennis, Steven Hazell, Costas Mitsopoulos, Colin Cooper, The ICGC-UK, TCGA Consortium, Kaitlyn P Russell, Paul Workman, Rosalind A Eeles, Bissan Al-Lazikani

**Author notes:** These authors contributed equally to this study (Stephanie T Schmidt, Zsofia Kote-Jarai, Adrian Larkeryd). Full member lists and their associated affiliations of these consortia are supplied in the two last tables of the Supplementary, Tables 4 and 5 (The ICGC-UK, TCGA Consortium).

## Abstract

Despite advances in understanding and treating Prostate Cancer (PCa), there has been little effort to systematically map the biology distinguishing Early-(EOPCa) and Late-(LOPCa) onset PCa. Around 25% of EOPCa cases present with metastatic spread or aggressive disease with earlier metastatic development. Some available lines of therapy are extending treatment trajectories and prolonging lives. However, there remains a critical clinical need to identify new therapeutic targets for EOPCa where life expectancy necessitates safer, more targeted treatment options. To our knowledge, here we present the largest systematic analysis of molecular profiles in EOPCa versus LOPCa, employing machine-learning-enabled algorithms to identify distinguishing biology and druggable targets for each age group. Distinct stromal signatures are uncovered in EOPCa, which are used to propose therapeutic opportunities herein. Moreover, our analysis identifies 50 druggable targets, 11 of which we confirm in PCa cell line genetic/pharmacological perturbation data. These findings provide the first specific, testable hypotheses in EOPCa, offering avenues for experimental validation and potential therapeutic exploitation, and, more generally, shed light on the intricate and distinguished molecular profile of this aggressive, poorly understood disease.

**One Sentence Summary:** Machine learning-enabled algorithms were utilized to identify distinguishing biology and associated druggable targets for early-onset prostate cancers.

## INTRODUCTION

Prostate cancer (PCa) is the most common solid cancer in men with greater than 1.4 million new cases annually diagnosed worldwide (*1*). Variability in age of onset and outcome presents a significant clinical challenge. There is some understanding of the evolution of mutational processes and the levels of heterogeneity at different stages of development, but very little is known about why some men develop clinically detected early-onset disease. Numerous studies focusing on identifying prognostic markers have been published (*2–4*), and key driver molecular events have been therapeutically exploited, particularly targeting AR regulation (abiraterone (*5*), enzalutamide (*6*)) and DNA damage repair pathways (PARP inhibitors (*7*) and immunotherapies (*8*)). The early onset of PCa (EOPCa), in particular, is an important clinical indication of higher probability of genetic predisposition. Some germline alterations (*BRCA2* (*9*)*, ATM* (*10*)) are known to be responsible for more aggressive outcomes in EOPCa, suggesting specific molecular trajectory/tumor progression and evolution. Although extensive data on the structure of PCa genomes have been published (*11–15*) and some evidence that EOPCa is associated with AR-driven homologous recombination defects has been reported (*16*), there has been no systematic analysis of molecular differences between EOPCa and late-onset PCa (LOPCa). Accordingly, no progress on differentiated, mechanism-driven treatment strategies for EOPCa have been presented. In general, EOPCa has been largely understudied.

In this study, we recruited 23 EOPCa patients for whole exome tumor sequencing, here referred to as ICR_BC. We combined these new data with our published data from TCGA_US (*12*) (https://www.cancer.gov/tcga) and ICGC_UK (https://dcc.icgc.org/) PCa studies to systematically map unique mechanistic signaling networks differentiating between EOPCa and LOPCa. This analysis provides new insights into the mechanisms of EOPCa through machine-learning based approaches and identifies novel molecular-based strategies for potential therapeutic targeting.

## RESULTS

The overall design of the study is shown in **Figure 1A**. In total, this study contains 688 participants, divided into two age cohorts to capture the signal of EOPCa versus LOPCa (**Figure 1B below and Table 1 in Methods**). The precise age cut-off for defining EOPCa varies in the literature and is between 50-65. To define the cutoff used in this work, we examined the age of onset distribution reported in the Surveillance, Epidemiology, and End Results ((SEER)*Explorer) database which includes PCa patient data from 1971-2023 (*17*). It shows an age of 60 as the inflection point in number of cases, which agrees with our data in this study as shown in **Figure 1B**. Therefore, EOPCa herein is defined as those diagnosed at 60 years of age or younger (n=301) and LOPCa as greater than 60 years (n=387). It is important to note that there are numerous factors that can additionally be used to group the patients such as Gleason scores and elevation of Androgen Receptor (AR). We find that Gleason scores of patients in the LOPCa cohort are typically higher than those in EOPCa (**Figure 1C**), but we do not find statistically significant difference in AR levels between the subgroups. For this analysis, we grouped patients solely on age cut-off and use the other factors as labels only.

**Figure 1.**
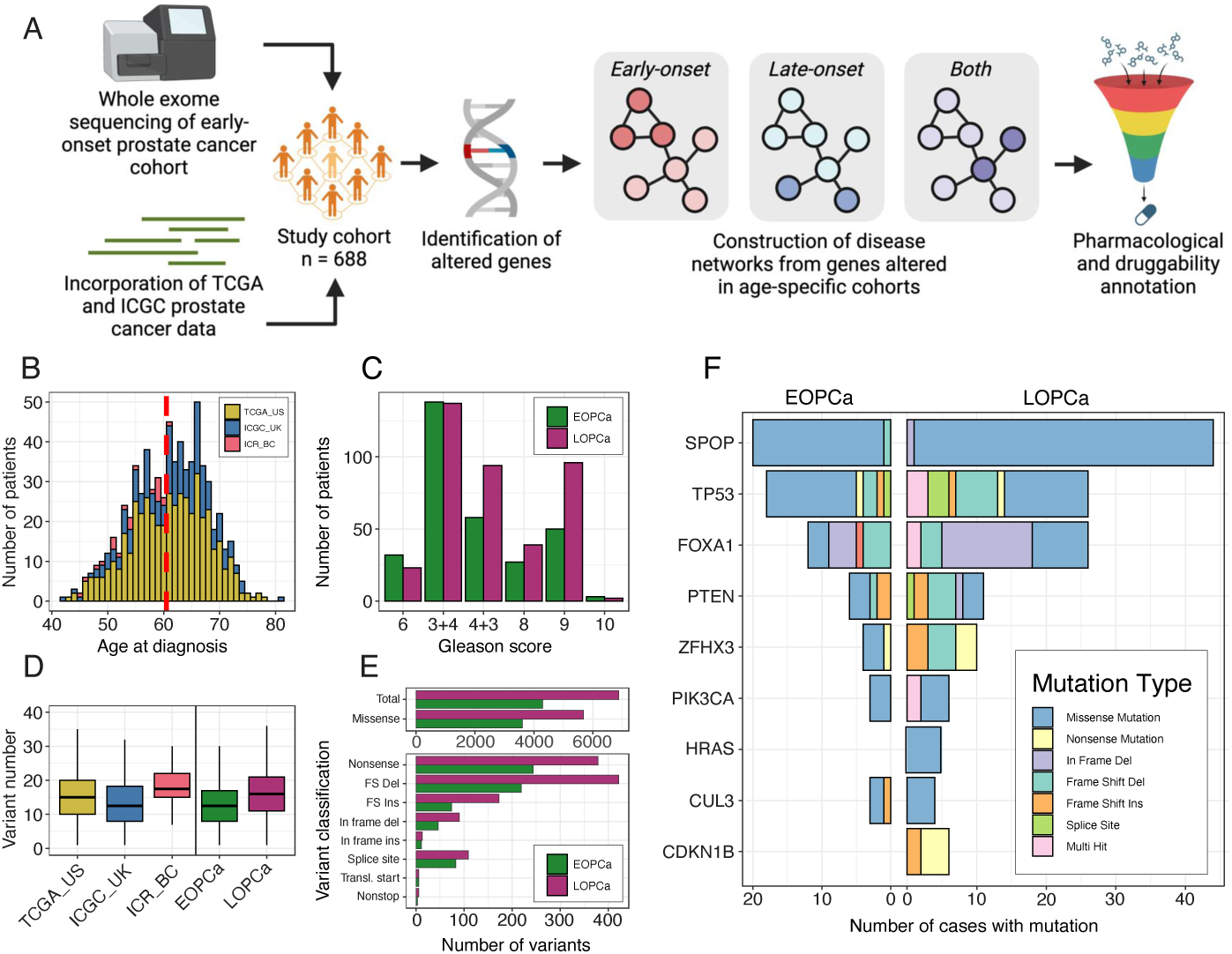
Workflow and descriptions of patient cohorts used to understand early versus late onset prostate cancer (E- or L-OPCa). **(A)** Study workflow (created with BioRender.com). **(B)** Age of onset histogram for all participants colored by study. Dotted red line shows age threshold by which participants were divided. **(C)** Gleason score for all samples colored by study. For samples with a total Gleason score of 7, primary and secondary division of 3+4 or 4+3 is shown. **(D)** Boxplot of total number of non-synonymous somatic variants by study as well as by selected age cohort. Boxplot outliers not shown. **(E)** Two graphs of variant classification colored by age cohort. **(F)** Co-bar plot of cases with mutations for genes identified by at least two methods. Mutations in EOPCa are shown at left and mutations in LOPCa at right.

**Table 1.** Patients recruited with PCa recruited from Royal Marsden Hospital for this study (ICR_BC) which we combined with available ICGC (ICGC_UK) and TCGA (TCGA_US) patient cohorts.

| Samples<br>Analyzed | Total | Early onset<br>(≤60 years) | Late onset<br>(>60 years) | NGS data |
| --- | --- | --- | --- | --- |
| ICGC_UK | 193 | 68 | 125 | Whole Genome |
| TCGA_US | 472 | 211 | 261 | Whole Exome |
| ICR_BC | 23 | 22 | 1 | Whole Exome |
| Total | 688 | 301 | 387 | Whole Exome +<br>Genome |

From this combined data, we first analyzed somatic alteration data including whole genome (n=193) and whole exome (n=495) data. We performed variant calling using the nf-core (*3*) pipeline Sarek (*4*) (version 2.7), with the tools Mutect2 (*5*) and ASCAT (*6*), and genome version GRCh38. Tumor Mutational Burden (TMB) between the three combined cohorts was largely similar although this appears lowest in ICGC_UK and highest in ICR_BC. We identified 4,294 non-synonymous somatic variants in 3,291 genes in EOPCa, and 6,875 variants in 4,738 genes in LOPCa. However, when comparing the EOPCa versus LOPCa cohorts, we find a statistically significant difference with higher TMB in LOPCa (Wilcoxon rank sum test, p-value = 3.22 x 10^-07^; **Figure 1D**). The higher number of mutations in LOPCa persisted across all mutation types (**Figure 1E**).

Given the relatively small amount of variant data in each age cohort, we applied three independent methods to determine recurrently mutated genes. First, an adapted use of Chang et al.’s method (*18*) was utilized to identify mutational hotspots (q-value <0.01). Secondly, MutSigCV was utilized to analyze the SNV data for additional significant hot spots (q-value<0.01). Lastly, we employed the ‘Cancer-Target Association’ (CTA) method from canSAR (*19*) which scores genes based on recurrent mutations in the gene while taking cohort burden and mutation impact into account. Considering all deployed methods, we find 31 genes with point mutations of which only nine are identified by at least two methods in the two cohorts. Mutations in recurring genes are shown in **Figure 1F**. For LOPCa, SPOP, FOXA1, PTEN, ZFHX3, HRAS, and CDKN1B genes all show higher mutation frequencies than in EOPCa. Of these, HRAS and CDKN1B mutations were unique to LOPCa. These mutation types were generally missense mutations regardless of subgroups, but LOPCa did exhibit other mutation types. The significance of these mutations is detailed further in the Discussion. Next, we identified ‘focal’ Copy Number Variations (CNVs) as being regions of chromosomes that have significant copy number alteration affecting five or fewer genes only (see Methods) and map these regions to 69 genes (as summarized in the study workflow in **Fig. S1**).

In order to construct the disease networks, we combined the mutated genes above with the 69 genes showing significant copy number alterations, resulting in 97 unique genes from the combined WES data. In total, for EOPCa, we identify 42 unique genes from amplifications, 18 from SNVs, 6 from deletions, 6 from both amplifications and deletions, and 1 from both an SNV and deletion. In LOPCa, 46 unique genes were altered, with 23 identified from deletions, 18 from SNVs, 4 from amplifications, and 1 from both an SNV and deletion. These sets form ‘seed genes’ that we use to construct the mechanistic networks.

We then applied our machine-learning-enabled workflow to generate protein interaction networks, which we previously applied to identify 22 mutated genes from prostate cancer patients (*20*). Briefly, the algorithm first identifies experimentally defined interactions between the seed gene using the canSAR interactome (*21*), which contains greater than one million experimentally determined protein-protein interactions. This typically results in a number of small, disconnected protein networks. We then iteratively connect these disconnected sub-networks algorithmically by identifying experimentally defined non-seed proteins that are (1) able to connect two or more subnetworks through interacting with their seed members and (2) demonstrate enrichment of their interactions with the seed proteins versus the rest of the human proteome. These additional proteins, which we term latent connectors, are not altered in patients but necessary for molecular communication between altered proteins. Finally, we perform the pharmacological annotation of the full network using up-to-date data from canSAR (*21, 22*) (https://cansar.ai) by labelling targets of approved drugs, investigational drugs, and chemical tools and those that have ‘druggable’ protein 3D structures. The resulting networks, shown in **Figure 2A-C**, contain 221 proteins. Fifty-eight proteins which are shared between EOPCa and LOPCa comprise 21 mutated or copy-number variant proteins and 37 latent connector proteins. Interestingly, using only simple bioinformatics on the same WES data, an AR mutation is lacking in this cohort and indicates that it is not a disease driver. However, our algorithm identified it as the central protein connecting the communication across all other proteins in the network, highlighting the ability of our methods to fill in key biological gaps in incomplete WES data.

**Figure 2.**
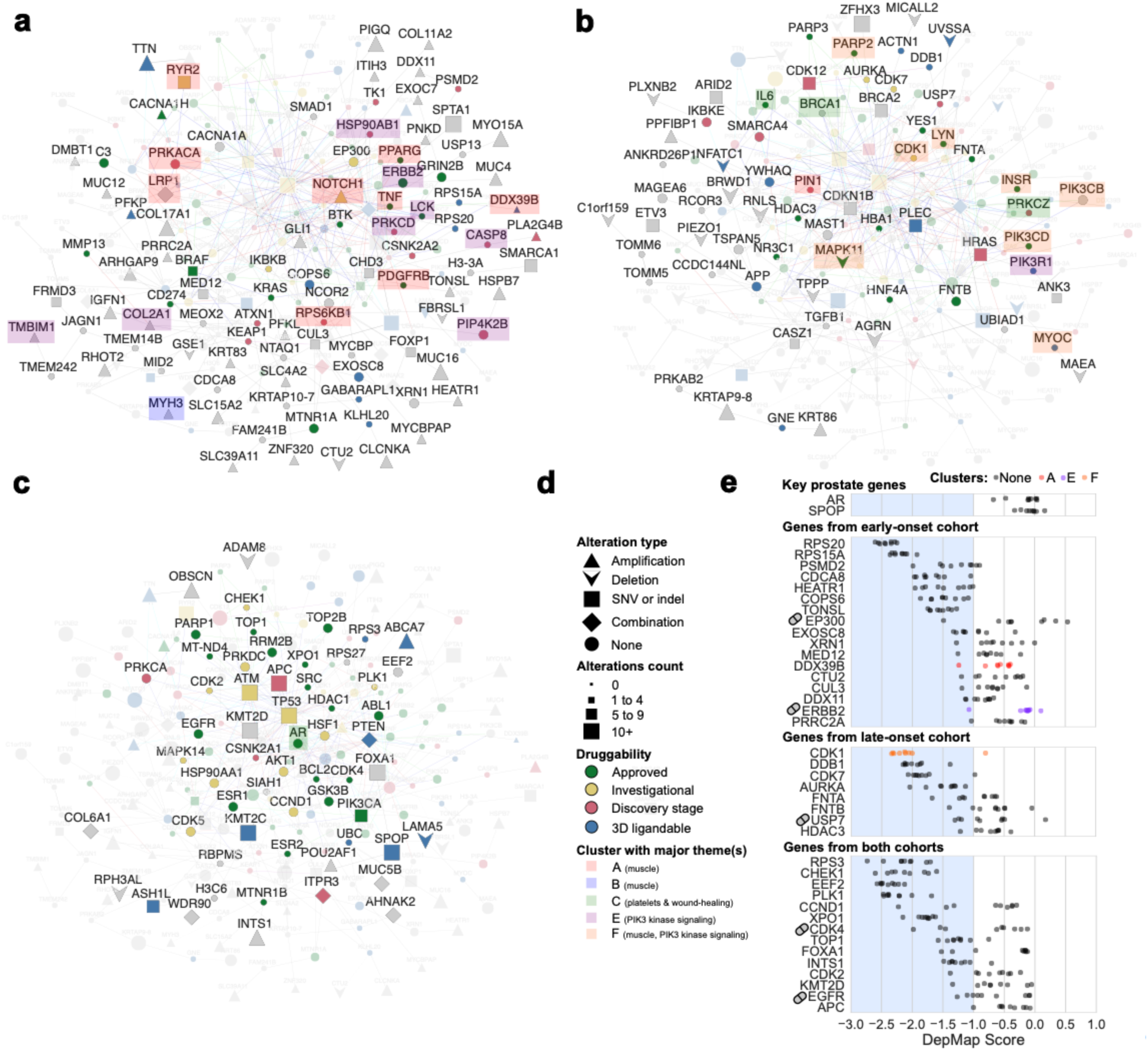
Full network of prostate cancer with annotation of pharmacology, druggability, and relevant pathways. Three views highlight each network subset: **(A)** Network of EOPCa emphasizing genes unique to it. **(B)** Network of LOPCa emphasizing genes unique to it. **(C)** Network emphasizing genes in both cohorts’ networks. **(D)** Legend for panels A-C. **(E)** Network genes observed to be a dependency (corresponds to blue shaded region) in at least one prostate cancer cell line from the DepMap project (*23*). Pill icons next to labels denote genes targeted by at least one drug/chemical tool in the Genomics of Drug Sensitivity in Cancer (GDSC) project(*24*).

To add further biological context around the WES-derived network, we used DEseq2 (*23*) to identify differentially expressed genes between EOPCa and LOPCa using the RNA-seq data available from the TCGA-PRAD study (178 EOPCa and 237 LOPCa samples). Genes with an adjusted p-value < 0.05 and absolute fold change > 1.5 were included in downstream analysis, along with the genes AR and SPOP, given their importance to prostate cancer (*25*). Our analysis returned 507 genes that were distinct between the two cohorts. We then constructed the expanded disease network seeded by the combined WES as well as differentially expressed genes **(Figures S2-S5**), and utilized Fisher’s Exact Test applied to the Gene Ontology (GO) Biological Process (BP) gene set collection(*24*) to perform the pathway annotation. These combined networks and their annotation provide a clear view of disease distinction as discussed subsequently.

From the pathway enrichment analysis of the distinct genes derived from each of the two disease networks (WES alone and WES + RNAseq, 663 genes in total), we highlighted pathways meeting the criterion of False Discovery Rate (FDR) < 0.05 and containing at least seven onset-age-specific genes in **Supplementary Figure S6**. Key drivers AR and SPOP were not differentially altered between the two subgroups, but we added them to the visualization for context. We annotated the differences between the early and late groups to highlight the distinct processes involved in each (**Figure 3** – truncated for ease of display; full figure in **Supplementary Figure S6**). It is important to remember that pathway names deposited into pathway databases and used by pathway enrichment methods often carry the original context of the experiments they were modeling when originally published. This requires us, as should be general practice, to examine the general patterns of pathways emerging rather than rely solely on specific names of pathways. As an example, our pathway analysis here includes ‘cardiac’ related pathways. Based on the other emerging pathways, these are more likely pointers to muscular or connective tissue signaling than specifically cardiac-related. We further discuss the presence of muscular signaling in EOPCa later in this work.

**Figure 3.**
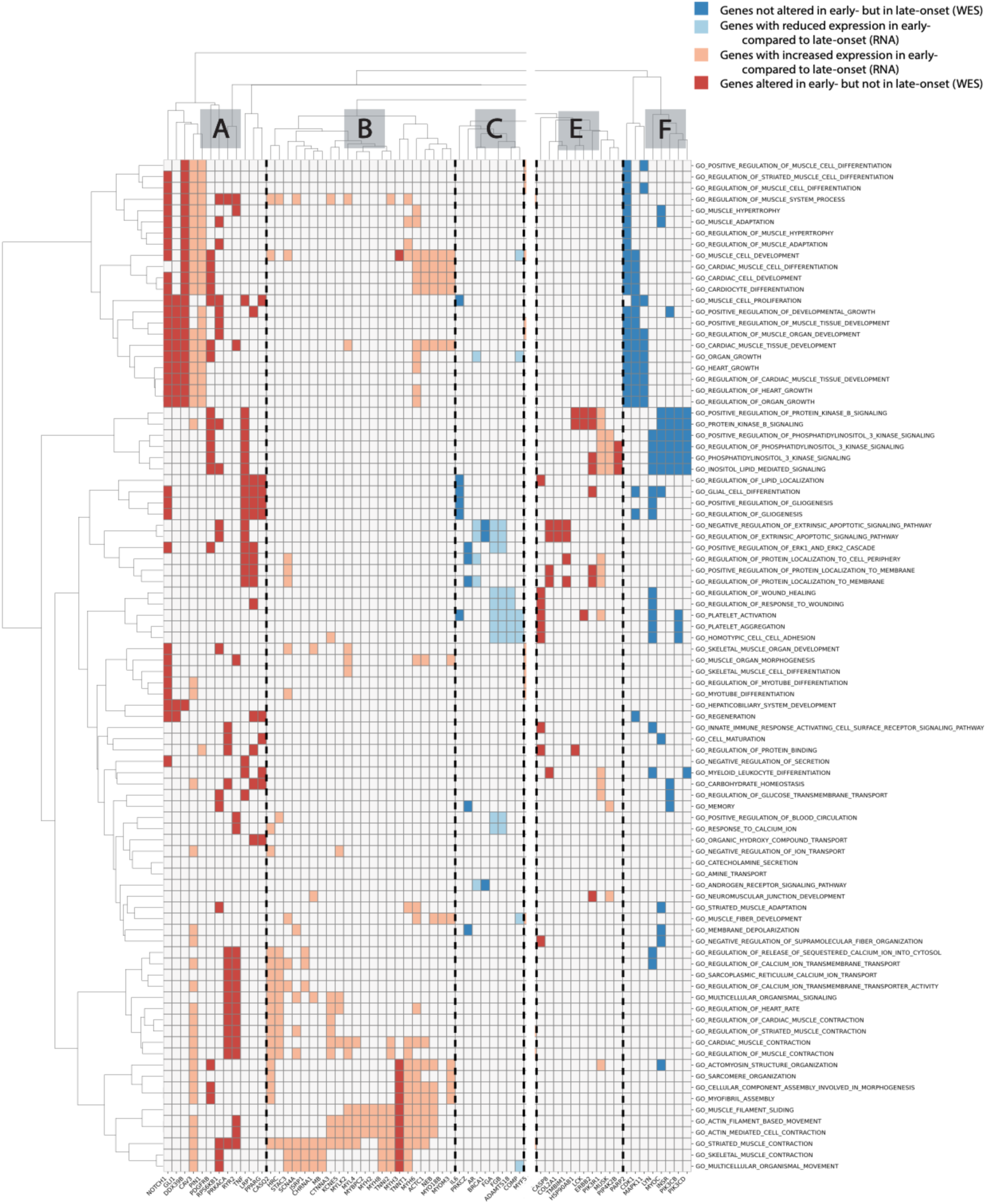
Pathway annotation of genes distinct in WES EOPCa versus LOPCa networks combined with differentially expressed genes between the two groups. X-axis lists the genes; Y-axis lists the significant pathways associated with them, colors indicate patient subgroups where a is gene is most important: red=EOPCa, blue=LOPCa. Full version of the figure is in **Supplementary** Figure 6.

The pathway analysis highlighted six major clusters of genes (groups A-F in **Fig. 2D and 3**) that differentiated between the early- and late-onset patients. Of these clusters, we observed that three (groups A, B, and F) included genes associated with muscle development and function, one with platelets and wound healing (group C), and two with PIK3 kinase signaling (groups E, F). Group D included diverse pathways without a clear overarching theme and therefore was not included in **Figure 2D**. The significant dysregulation of groups A, B, and F points to differences in the stroma and muscular microenvironment between EOPCa and LOPCa tumors. The genes of the platelets and wound healing group (group C) were less altered or expressed in the early onset cohort. Finally, genes associated with PIK3 kinase signaling also demonstrated significant dysregulation between the two onset-age groups and included some genes more (group E) or less (group F) altered or expressed in the early onset cohort. The link between PIK3 and the mTOR pathway with prostate cancer is becoming an area of increasing interest (*26, 27*). We have shown previously that germline alterations in the PI3K/AKT/mTOR pathway associate with biochemical recurrence (*22*) and also note that >40 clinical trials are underway for PCa with PTEN loss (https://clinicaltrials.gov; **Table S1**). PTEN was previously identified as a frequent mutation in EOPCa by Gerhauser et al. along with several other age-related genomic alterations and a clock-like enzymatic-driven mutational process (*16*). We compare these findings further in the Discussion.

The muscle-related signal is robust and intriguing. To verify that this muscle signal is not merely an artifact of the age of the EOPCa cohorts, we examined this pattern in patients with other solid tumors in the same age range and found that this muscle-related signal is indeed specific to PCa. For example, in melanoma, we found that 368 genes were differentially expressed between the two age groups, but no pathways (FDR < 0.05) were found to be enriched amongst these genes. As a further control, we performed similar analysis in glioblastoma (GBM) and found that 82 genes were differentially expressed between the two age groups. These genes were associated with five pathways (FDR < 0.05), namely cell fate specification, innervation, nerve development, negative regulation of neuron apoptotic process, and neuron apoptotic process. No significant associations with muscle-related pathways were found in GBM. These data suggest that the muscle-related signals observed in prostate cancer are not an artifact of the younger age of the EOPCa cohort but instead reflect the mechanisms of the disease itself and warrant future validation.

The integrative analysis so far has identified altered genes, latent connector proteins, and mapped key pathways onto these. To verify whether and which genes in the disease network have biologically functional impact, we interrogated the DepMap (*23*) database which contains genome-wide CRISPR screens against nine PCa cell lines. Thirty-nine genes identified by our analyses were confirmed genetic dependencies in at least one of the nine PCa cell lines profiled by CRISPR knock-out screening in DepMap (**Figure 2E**). Of the 39 genes verified to be dependencies in at least one PCa cell line in DepMap, 17 dependencies were uniquely from the EOPCa-network (e.g., EP300, DDX11, and RPS20). Eight were identified from the LOPCa-network (e.g., AURKA, CDK1, and CDK7), and 14 genes were shared between EOPCa and LOPCa (e.g., APC, CDK2, CDK4, and CHEK1). From the nine PCa cell lines tested in DepMap, we determined that three – WEP1NA22, LNCAPCLONEFGC, and VCAP – were originally derived from patients <60 years old (**Table S2**). Twenty-six of the dependencies identified from EOPCa were indeed dependencies in these three lines (**Table S3**). These include EP300, CHEK1, and APC. To ensure that our resulting confirmed dependencies are significant, we performed 1000 statistical trials of randomly selected genes of the same list size and observed that the chance of observing the same results was less than 0.001% (**Supplementary note 1, Fig. S7**).

We then queried the Genomics of Drug Sensitivity in Cancer (GDSC) database (*24*) for pharmacological perturbation data to complement the genetic perturbation information provided by DepMap. Of the proteins identified from our networks, 36 had drugs tested in PCa cell lines. In total, 16 targets had drugs that showed activity in at least one PCa cell line, defined as a reported z-score of the natural logarithm of IC50 less than -1.5. Four were from the EOPCa-network (KRAS, EP300, ERBB2, BTK), five from the LOPCa-network (TGFB1, PIK3CD, PIK3R1, USP7, PIK3CB), and seven from both cohorts (PIK3CA, AKT1, ESR1, ESR2, EGFR, CDK4, GSK3B).

In total, our disease networks identified 221 proteins in our PCa networks. Of these, 97 were significantly altered in one or both cohorts. Of these, 13 were confirmed as dependencies in PCa cell lines using genetic/drug perturbation. A further 124 were identified by our algorithm to be important for disease communication although not significantly altered in either patient cohort. Of these, 37 were similarly confirmed by genetic and/or drug perturbation in PCa cell lines. Of the confirmed dependencies that are significantly altered in patients, eight are derived uniquely from the EOPCa-network and four were derived from the shared network.

## DISCUSSION

Our work identified the dependence of several genes in both EOPCa and LOPCa such as PTEN and TP53, which as mentioned were identified previously by Gerhauser et al (*16*). PTEN was found to have the highest rate of biallelic inactivation, followed by TP53, specifically in EOPCa cohorts. TP53 was the most altered by nonsynonymous SNVs in EOPCa cohorts.

Additionally, some of the APOBEC-associated SV breakpoints, which include PTEN, indicated that APOBEC mutations, an age-associated mutational process, are likely responsible for early PC mutations. Both PTEN and TP53 were also found to be present in aggressive PCa molecular subtypes. Interestingly, another APOBEC-associated gene, BRCA2, was found in earlier disease progression, whereas we only identified BRCA2 in the LOPCa cohorts. The study, while including some of the clock-like APOBEC mutagenesis genes from Gerhauser et al. (*16*), largely implicate genes involved in muscle development/function, PIK3 kinase signaling, and platelets/wound healing. Genes associated with muscle development and function which were enriched in EOPCa could indicate high EMT activity and/or smooth muscle contamination within primary tumors, although experimental exploration would be needed to confirm this.

The systematic, high-throughput genetic perturbation data provided by DepMap^1^ identifies genes required for cell growth, comprising an invaluable resource for understanding genetic dependencies and providing initial confirmation of our findings. Further validation, however, in patient-relevant models will be required to progress any such findings to the clinic given the following caveats. Firstly, existing 2D-PCa cell line models have known limitations (*28*); the standardized assays and conditions in DepMap, necessary for the systematic large-scale nature of the resource, could lead to findings which may or may not actually be significant in PCa disease. For example, we expect that beyond the thirty-nine genes that were dependencies in the prostate cancer cell lines profiled by DepMap, there may be additional dependencies at play in prostate cancer which could be missed. This is the case for the key PCa driver gene, AR, which is not a dependency in prostate cancer by DepMap with an average Chronos score of - 0.16. The missed dependency can in part be explained by the fact that prostate cancer cell lines are particularly challenging to grow (*29*). The implications are that dependencies will be more difficult to capture in slow growing cell lines within the time scale of the DepMap standardized assay (∼72 hours), which is optimized for the ‘average’ cell line. The second caveat is that prostate cancer cell lines, like any model system, are not necessarily representative of the disease itself. We reviewed the prostate cancer cell lines included in DepMap and found that their origins were varied, including a serially propagated tumor xenograft, a normal prostate cell line, and a benign prostatic hyperplasia, in addition to the expected patient tumor samples (*2*) (**Table S2**). Despite the limitations which we have described here for completeness, public databases like DepMap provide an invaluable, large-scale experimental dataset for contextualization of our findings and strong rationale for deeper experimental validation.

Although PCa cell lines are non-ideal models, this initial confirmation demonstrates the potential to identify druggable/actionable targets for each of the cohorts. Interestingly, in addition to the 65 drug targets with FDA-approved or clinical stage drugs (colored in green and yellow in the networks in **Figure 2A-C**), our analysis identifies a further 50 druggable targets (red and blue in **Figure 2A-C**). Of these, 17 came from the 97 altered genes, and 11 were found to be genetic dependencies in PCa cell lines. Six of these PSMD2, COPS6, RPS15A, EXOSC8, DDX39B, RPS20) are derived from the EOPCa cohort and can potentially be exploited for the discovery of novel therapeutics. While the exact functional relevance of these druggable targets is unknown, they are linked to established molecular processes in prostate cancer(*30–35*) and thus are sensible to evaluate further. Because PCa cell lines do not represent the complexity of patients nor the tumor microenvironment, we anticipate greater confirmation and insight to come from validation experiments in biological models which better represent patients. Regardless, these early signatures in PCa demonstrate the potential therapeutic possibilities available to these patients.

This, to our knowledge, is the first systematic analysis of the differences in the underlying mechanisms differentiating EOPCa versus LOPCa. We utilized data from 688 patients and applied our machine-learning-enabled approach to identify distinguishing mechanisms and druggable targets for each cohort. We confirmed 11 druggable targets as molecular dependencies in PCa cell lines using genetic and drug perturbation data and highlighted six novel potentially druggable targets for further examination in EOPCa. The stromal component is the most distinguishing feature between EOPCa and LOPCa. This, for the first time, provides testable biological hypotheses unique to EOPCa that can, after validation in appropriate models, be potentially exploited for therapeutic intervention.

## MATERIALS AND METHODS

### Experimental Study Design

A total of 23 men with EOPCa prostate cancer were recruited from The Royal Marsden Hospital to understand differences between EOPCa and LOPCa. These 23 patients were combined with available data from both the International Cancer Genome Consortium (ICGC) and The Cancer Genome Atlas (TCGA) as described in **Table 1** to provide a more complete dataset. The distinction of EOPCa vs LOPCa varies in literature, so we defined our age criteria for EOPCa as being less than or equal to the peak of the age-onset distribution published by SEER*Explorer (*17*), which was 60 years of age. Within our dataset of 23 patients, there were no patient outliers. Additionally, all patients were recruited given relevant ethical regulations. Ethical approval was obtained from the respective local ethics committees and from The Trent Multicentre Research Ethics Committee. All patients were consented to ICGC standards.

From the combined data in **Table 1**, we expected to highlight distinguishing gene signatures and networks between EOPCa and LOPCa. Further, we hoped to identify a number of exploitable drug targets. Upon investigation, our pathway analyses (detailed below) identified several distinct genes in EOPCa which regulate muscle connectivity. We suspected that the presence of these genes was true and not simply an artifact of the younger age of the EOPCa cohort, which we explored in this paper using methods described in the *“Examination of muscle-related signal in other solid tumors” Methods*.

### Sequencing of ICR_BC samples

Germline DNA samples from the 23 ICR_BC PCa cases (Gleason ≥7, tumor stage 2a-3b) were extracted from whole blood. Tumor DNA was extracted following radical prostatectomy and prior to additional treatment. 22 cases were diagnosed with PCa aged 60 or under (mean=54), and one case was diagnosed at 61.

DNA samples were fragmented using a Covaris E220 Ultrasonicator and exome sequences enriched using Agilent SureSelect XT2 Human All Exon V5 baits, in pools of 8 samples with 7 bp molecular barcodes. Pools were sequenced on an Illumina HiSeq 2500 instrument (v4 chemistry, 2 × 100 bp reads). Samples were assessed for sufficient coverage of >80% of bases at ≥20× sequencing depth (genomic samples) or ≥100× sequencing depth (tumor samples). As mentioned above and as previously described (*2*), these data were combined with prostate cancer sequencing data from the IGGC (ICGC_UK; https://icgc.org/) and TCGA (TCGA_US; https://www.cancer.gov/tcga)^1^.

### Variant calling

Variant calling was performed using the nf-core (*3*) pipeline Sarek (*4*) (version 2.7), with the tools Mutect2 (*5*) and ASCAT (*6*), and genome version GRCh38. Mutect2 analysis was performed with a panel of normals respective to each data set and with the argument *--dont-use-soft-clipped-bases true*. A panel of normals for Mutect2 was created for each of the three datasets in the study, using GATK (*5*) (version 4.1.3.0). For the TCGA_US dataset, a random selection of 100 of the normal samples were included in the panel of normals. For the ICGC_UK dataset, 243 normal samples from the same sequencing run were used to create the panel of normal. For the ICR_BC dataset, all 23 normal samples were included in the panel of normals.

### Filtering of SNVs

The exome capture regions of TCGA_US and ICR_BC were intersected to create a set of common regions captured for all samples (*7*). Variants called by Mutect2 from all samples, including the WGS dataset ICGC_UK, were filtered to include only variants in these regions. Further, the variants were filtered to keep only variants with the PASS flag from Mutect2, as well as a total depth of ≥25 and alt allele depth of ≥10. The remaining variants were annotated with VEP (*36*) (version 104) and converted to MAF format using vcf2maf (*9*). Four samples were removed due to a somatic mutation rate >10/Mb.

### SNV cancer target association & Recurrent Gene Analysis

To identify mutational hotspots (i.e., recurrent genes), the two cohorts were analyzed with an adapted method of the one developed by Chang et al. (*18*). Chang et al.’s method was previously utilized to identify mutational specificities in prostate cancer in 2016; our analysis adapted an updated version of the reference genome, with the R package BSgenome.Hsapiens.UCSC.hg38 (*11*). Genes containing a hotspot with a q-value <0.01 were considered significant hits. The SNV data was also analyzed using MutSigCV (*12*) (version 1.41) in MATLAB R2020b and previously used to identify biomarkers in radiation toxicity for prostate cancer. Genes with a q-value <0.01 were considered significant hits. A third complementary method utilizing canSAR-CTA was implemented to calculate associations between genes and each cohort. CanSAR-CTA scores genes based on recurrent mutations in the gene while taking cohort burden and mutation impact into account as previously demonstrated for EGFR (*19*). Mutation impact was annotated using snpEff (*13*) and with snpSift (*14*) Meta_SVM score and phastCons100way_vertebrate from dbNSFP (*15*). The CanSAR-CTA method results in a continuous score, of which the top 20 genes were added to the significant hits from the other two methods.

### Copy number variants

The segments produced by ASCAT (*6*) were annotated with gene locations with reference to GRCh38, and segments overlapping chr6:29,723,434-33,409,896 were removed to reduce complexity due to the HLA genes. GISTIC2 (*20*) (version 2.0.23) was used to analyze the ASCAT segments from two cohorts. The regions identified by GISTIC2 with a q-value <0.01 were selected as regions which might contain driver genes. To reduce the introduction of noise by passenger genes in large copy number events, only regions containing five or fewer genes were selected. Additionally, each individual sample was investigated in the regions identified by GISTIC2 by looking at fold change against the chromosome arm median. A fold change of 2.2 was used to call a deletion or amplification. If the median absolute copy number of the arm was zero, no amplifications or deletions were called on that arm. Only regions which contained amplified or deleted segments in at least 1% of patients were kept. All genes located in these focal copy number variant rich regions were added to the genes identified with the SNV data.

### Protein interaction networks

We constructed the disease networks using an updated version of the method previously described by our work (*20*). First, we defined a seed gene list as the combined altered list of genes from both the early- and late-onset cohorts. We generated two seed lists: one with only the alterations identified from Whole Exome Sequencing (WES) analysis and the second from the combined WES and differential gene expression data; the number of alterations between the two groups are summarized in the Supplementary **(Fig. S8).**

We then retrieved all experimentally determined protein-protein interactions between members of the seed list from the canSAR interactome (*21*), meaning that local protein interactions determine the link between seed genes. This yielded a small number of connected networks, but a number of seed proteins remained disconnected. We then applied an iterative algorithm to identify additional proteins (nodes) from the interactome that can help connect the disconnected proteins as well as the different small networks into a single connected large network (*2*). Some genes used in this seed list may or may not produce a protein depending on their own and mutations and those upstream, although we applied additional filtering which includes only the most enriched genes (and, presumably, protein-translation) as described subsequently.

For each added node, we calculated first-neighbor enrichment as the ratio of the number of edges connecting a node to seed nodes, over the total number of edges connecting that node to other nodes in the interactome. First-neighbor enrichment was then used to decide which transcriptional and complex edgesto keep, retaining only the nodes with the highest enrichment as defined by their interaction in the wider interactome. Additionally, any separate subnets which did not contain any of the seed nodes were removed from the graph. First-neighbor enrichment was again calculated against the seed nodes in the network and shortest paths between all seed nodes were determined. Hit enrichment in the calculated shortest paths was evaluated and the best candidates for each pair of seeds were selected based on the highest hit enrichment. Nodes which were not present in any of the enriched shortest paths were removed from the graph. To shrink the network further, all non-articulation, non-seed nodes were removed. Approved and investigational drug targets as well as earlier discovery stage compounds and 3D ligandable targets connected to at least two seeds were added to the network. Graph simplification continued by removing drug-drug edges that were not bridges, and any prior articulation points that were no longer required, provided they were not druggable or seeds. Quality probes and 3D-ligandable targets that were not seeds or articulation points were removed to produce the final network.

### Expression analysis from TCGA-PRAD Study

Genes present in the network of only one age group were included in pathway analysis to understand differences between them. To complement these genes, additional genes unique to each age group were identified from the TCGA_US cohort by performing differential expression analysis via DESeq2 (*23*). DESeq2 is routinely utilized to identify differentially expressed genes within different cohorts and has been employed by other prostate cancer studies (*37*). Genes with an adjusted p-value < 0.05 and absolute fold change > 1.5 were included in the downstream analysis, along with the genes AR and SPOP, given their importance to prostate cancer (*25*).

Pathway annotation was carried out using a generalized approach based on Fisher’s Exact Test and applied to the Gene Ontology (GO) Biological Process (BP) gene set collection (*24*). Gene sets meeting the criterion of FDR < 0.05 and that also had an overlap of at least seven genes were visualized in a heatmap to highlight the presence or absence of each gene within each pathway. The same approaches for differential expression analysis and pathway annotation were applied to RNA-seq data from primary tumors for patients under and over 60 years of age in the TCGA_GBM and TCGA_SKCM projects (https://www.cancer.gov/tcga) to examine muscle-related signal (discussed subsequently in *Examination of muscle-related signal in other solid tumors* section of *Materials and Methods*).

### Validation in public datasets

Genes from the network constructed were contextualized in DepMap (*38*) (Public 22Q4 release) and the GDSC (*24*) (GDSC2 release) databases of genetic and pharmacological perturbation, respectively. Genes with a reported DepMap score of < -1 were considered genetic dependencies from DepMap, while targets with a reported z-score of the natural logarithm of IC50 < -1.5 were considered to be sensitivities.

### Examination of muscle-related signal in other solid tumors

We observed muscle-related signals as a major theme returned by pathway enrichment analysis in comparing genes distinct to early-onset prostate cancer (EOPCa) and late-onset prostate cancer (LOPCa) that were either differentially altered or expressed. To ensure that this signal is not an artifact due simply to the younger age of the EOPCa cohort, we examined whether such a muscle signal would emerge in other solid tumors when cohorts are split into the same age groups. We performed the analysis as follows:

1. We gathered RNA-seq data available from primary tumors in male patients from the TCGA_GBM and TCGA_SKCM projects (https://www.cancer.gov/tcga).
2. We selected glioblastoma because there is no muscle in the vicinity of these tumors to contribute to a muscle-related signal. This acted as a negative control to ensure that the algorithm did not erroneously introduce a muscle signal. In contrast, we selected malignant melanoma because of muscle tissue proximal to these tumors. Thus, if the muscle signature observed in prostate cancer (PCa) was due to simply age difference, we expected the muscle signature to emerge in this melanoma cohort.
3. We then confirmed that the cutoff of 60 years to separate the patients with available data provided comparably sized groups for both glioblastoma (49 patients <60 years old, 50 patients >60 years old) and melanoma (24 patients <60 years old, 37 patients >60 years old).
4. We conducted differential expression analysis for these two groups in each indication followed by pathway enrichment analysis as we described for the PCa analysis in above *Methods* section.

### Statistical Analysis

Our work employed several statistical tools (e.g., pathway enrichment, protein interaction network, etc.) as described in detail above. These analyses and their appropriate criteria for replicating our methods are summarized in Table 2 within the *Tables*.

**Table 2.**
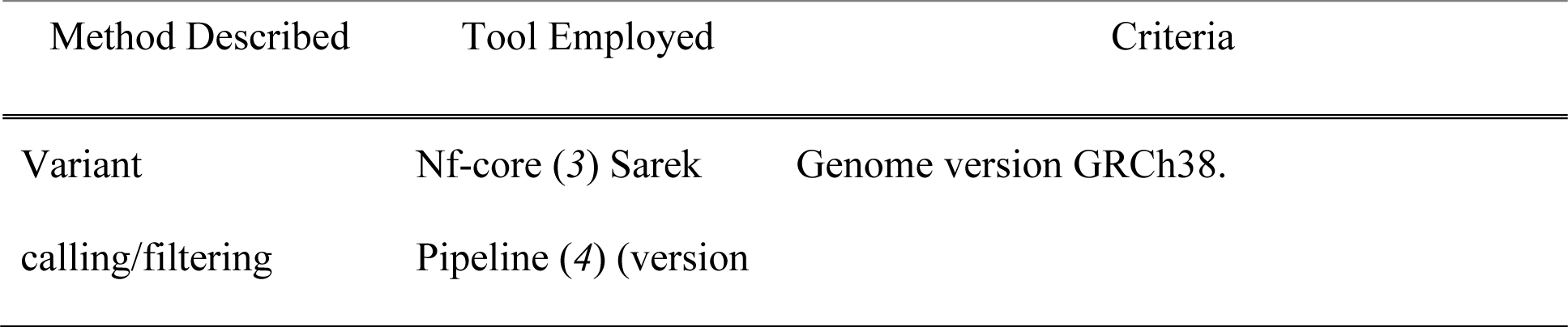
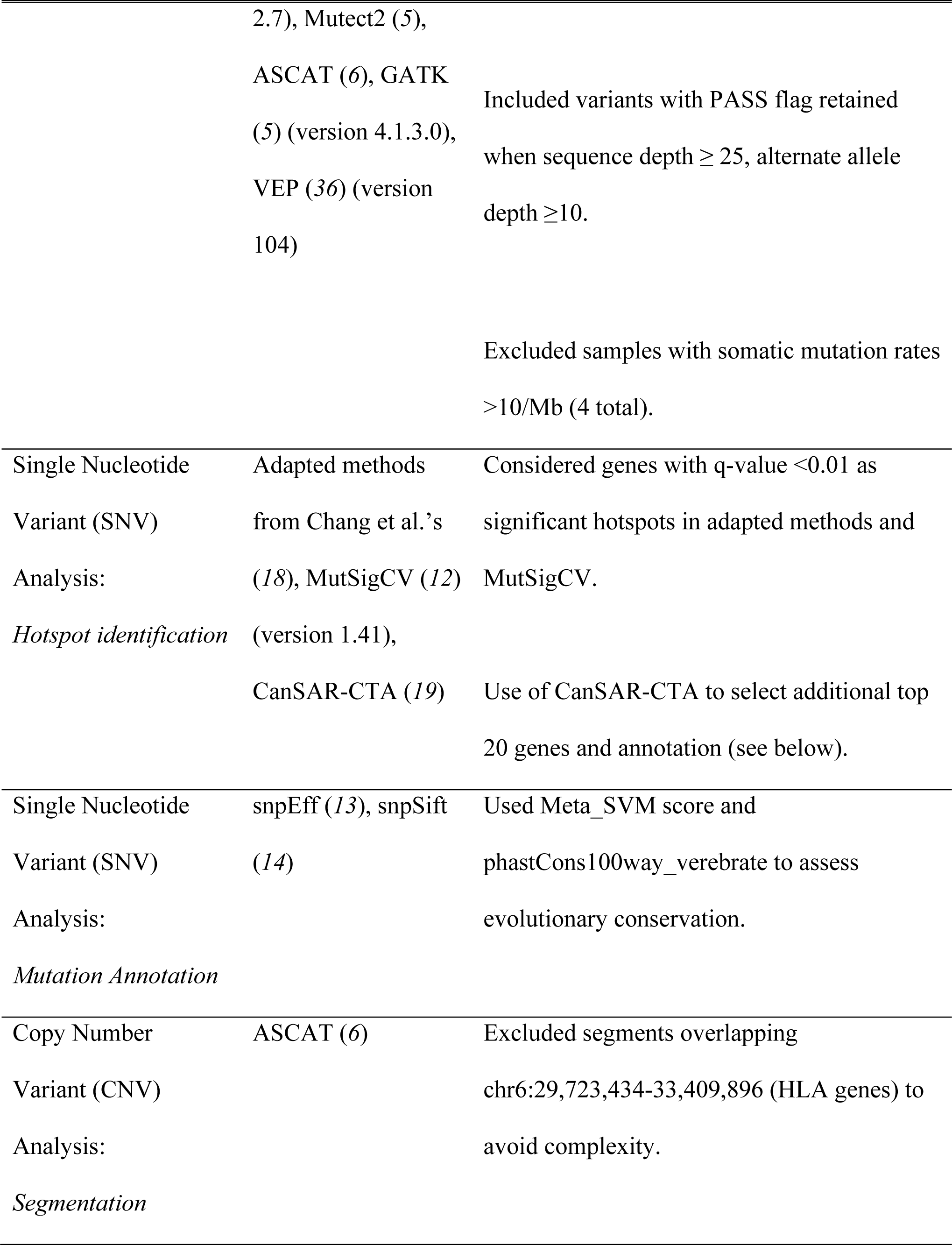
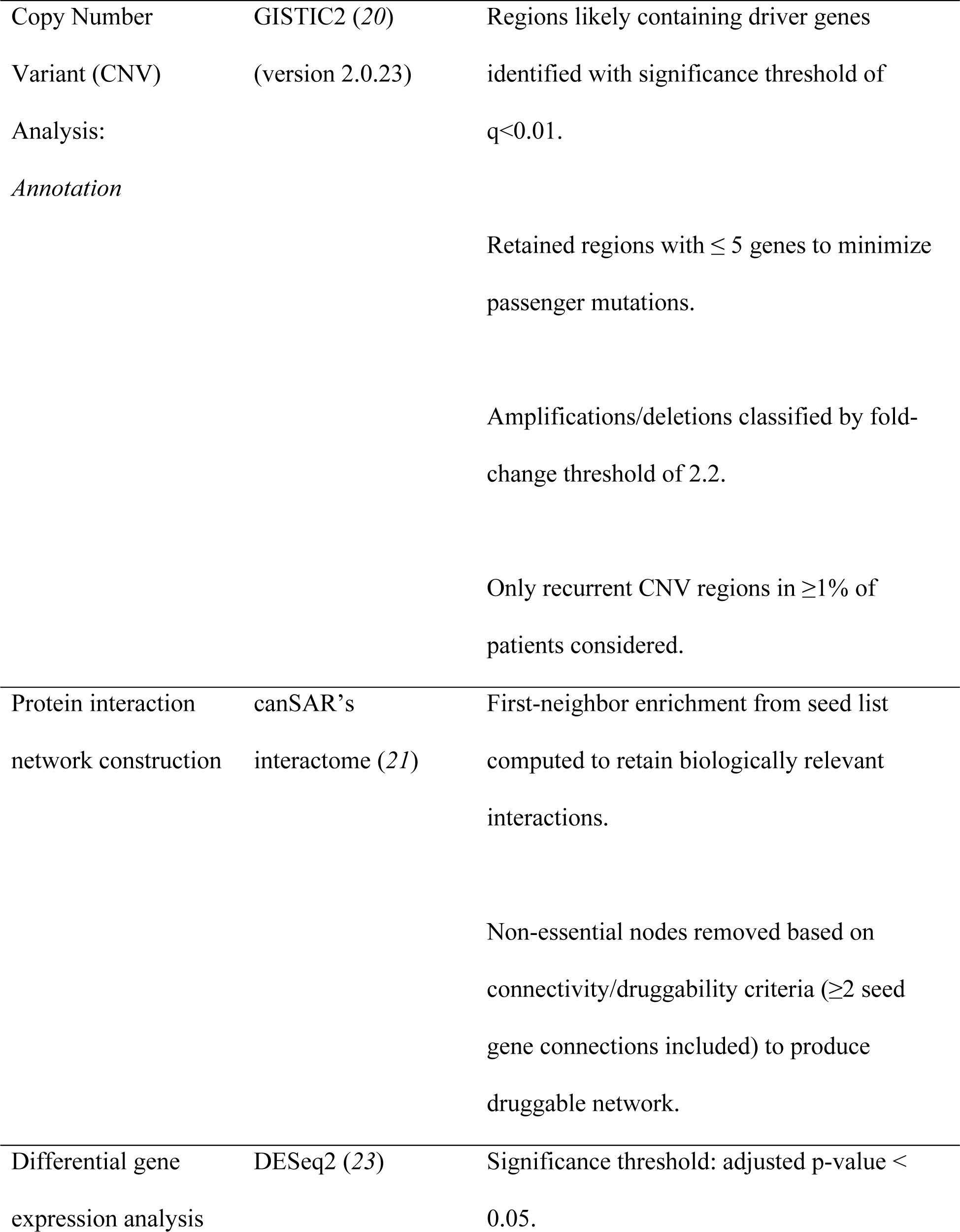
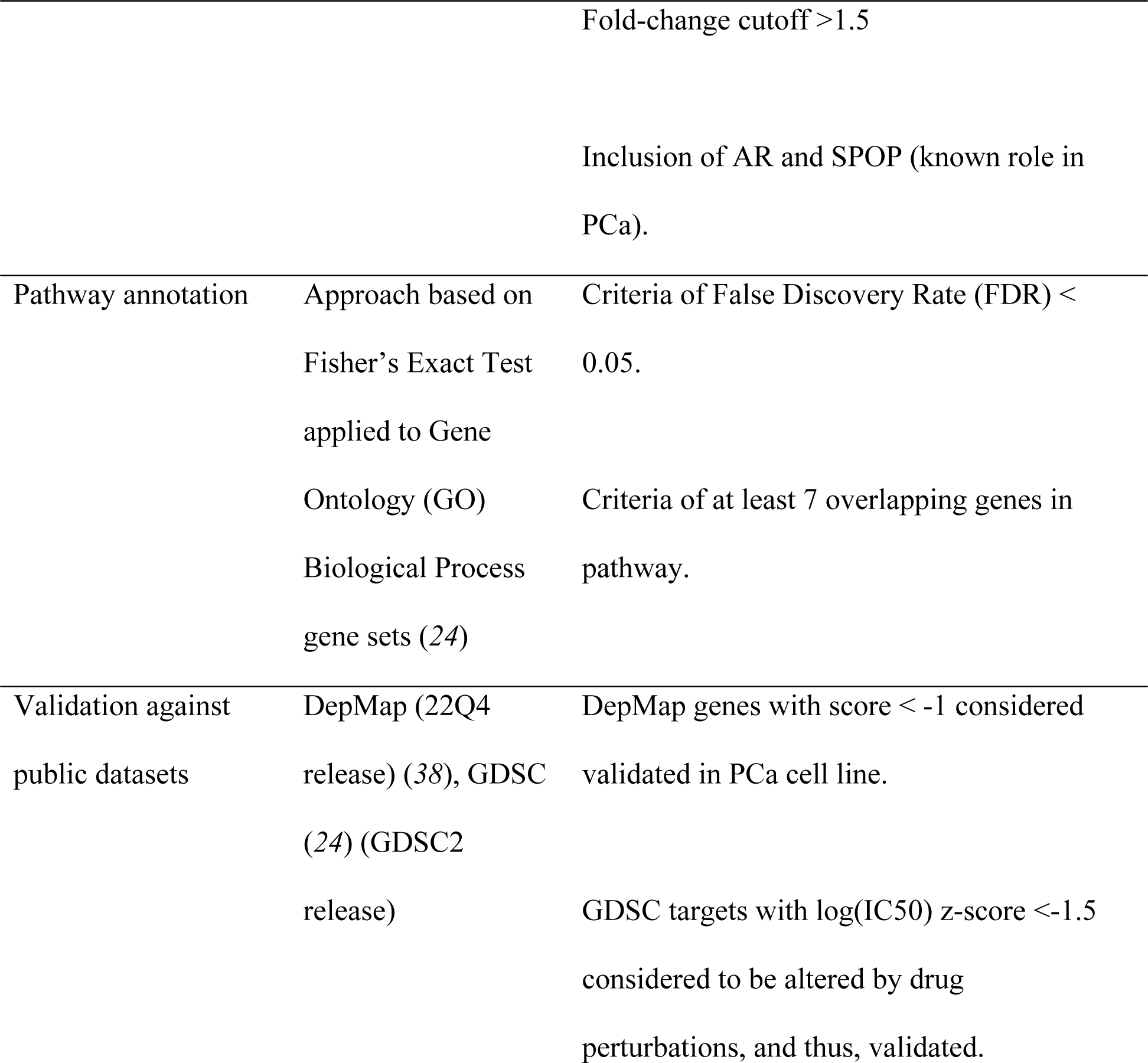
Summary of methods employed within this work, the tools to execute them, and the criteria used in each tool.

## List of Supplementary Materials

Figure S1: Expanded study workflow

Figure S2: Network of differential WES And RNA-seq genes annotated by pathway cluster: EOPCa

Figure S3: Network of differential WES And RNA-seq genes annotated by pathway cluster: LOPCa

Figure S4: Network of differential WES And RNA-seq genes annotated by druggability: EOPCa Figure S5: Network of differential WES And RNA-seq genes annotated by druggability: LOPCa

Figure S6: Pathway annotation of differential WES network genes and differentially expressed genes

Figure S7: Number of genetic dependencies identified using a gene set of the same size

Figure S8: Venn diagram of genes identified from WES alterations, identified by differential expression, and included in the main networks

Supplementary Note 1: Significance of the number of dependencies identified

Table S1: Interventional clinical trials from Clinicaltrials.Gov for PCa with patients harboring PTEN loss

Table S2: Description of 9 PCa cell lines with CRISPR profiling data available

Table S3: Dependencies occurring in EOPCa observed in cell lines derived from EOPCa Samples

Table S4: TCGA Consortium Members Table S5: ICGC Consortium Members

## Acknowledgments

The authors thank those men with prostate cancer and the subjects who have donated their time and their samples to enable this research. BA-L is a Cancer Prevention & Research Institute of Texas (CPRIT) Scholar in Cancer Research. PW is a Cancer Research Life Fellow. We also acknowledge support from the Bob Champion Cancer Trust and the DJ Fielding Medical Research Trust. BA-L acknowledges funding from Cancer Prevention and Research Institute of Texas (CPRIT) Established Investigator Award (RR210007), The Commonwealth Foundation, The Lyda Hill Foundation and the CRUK Drug Discovery Committee strategic award for canSAR. We acknowledge support from Cancer Research UK C5047/A14835/A22530/A17528, Prostate Cancer UK, the NIHR (National Institute of Health Research) support to the Biomedical Research Centre at the Institute of Cancer Research and the Royal Marsden NHS Foundation Trust.

## Funding

Bob Champion Cancer Trust (BA-L)

DJ Fielding Medical Research Trust (RAE)

Cancer Prevention and Research Institute of Texas (CPRIT) Established Investigator Award RR210007, (BA-L)

The Commonwealth Foundation, (BA-L)

The Lyda Hill Foundation, (BA-L)

CRUK Drug Discovery Committee strategic award, (BA-L) Cancer Research UK C5047/A14835/A22530/ A17528, (RAE) Prostate Cancer UK, (BA-L)

NIHR (National Institute of Health Research) support to the Biomedical Research Centre at the Institute of Cancer Research, (BA-L)

Royal Marsden NHS Foundation Trust, (BA-L)

CRUK Programme Grants [C309/A31322 and C309/A11566], (PW) Strategic Award, Infrastructure Award [C309/A27413], (PW)

CRUK Childrens’ Brain Tumor Centre of Excellence [C9685/A26398/RG93685], (PW) CRIS Cancer, (PW)

The Institute of Cancer Research, (PW) Chordoma Foundation, (PW)

Mark Foundation, (PW)

Bone Cancer Research Trust, (PW)

## Author contributions

Each author’s contributions are listed below following CRediT guidelines:

Conceptualization: BA-L, RAE Methodology: BA-L, RAE

Investigation: ZK-J, QK, SM, ND, SH, STS, AL, JC, BO Visualization: ZK-J, QK, SM, ND, SH, STS, AL, JC, BO

Funding acquisition: BA-L, RAE

Project administration: BA-L, RAE Supervision: BA-L, RAE

Writing – original draft: STS, AL, ZK-J, PW, RE, BA-L Writing – review & editing: All authors

## Data and materials availability

The ICGC data are available from the ICGC consortium under standard request protocols: https://docs.icgc.org/download/data-access. The ICR_BC data can be requested by contacting.

Code is available at our team’s GitHub: https://github.com/a3d3a.

## References and Notes

1. H. Sung, J. Ferlay, R. L. Siegel, M. Laversanne, I. Soerjomataram, A. Jemal, F. Bray, Global Cancer Statistics 2020: GLOBOCAN Estimates of Incidence and Mortality Worldwide for 36 Cancers in 185 Countries. CA: A Cancer Journal for Clinicians 71, 209–249 (2021).

2. P. J. Boström, A. S. Bjartell, J. W. F. Catto, S. E. Eggener, H. Lilja, S. Loeb, J. Schalken, T. Schlomm, M. R. Cooperberg, Genomic Predictors of Outcome in Prostate Cancer. European Urology 68, 1033–1044 (2015).

3. J. Cuzick, G. P. Swanson, G. Fisher, A. R. Brothman, D. M. Berney, J. E. Reid, D. Mesher, V. O. Speights, E. Stankiewicz, C. S. Foster, H. Møller, P. Scardino, J. D. Warren, J. Park, A. Younus, D. D. Flake, II, S. Wagner, A. Gutin, J. S. Lanchbury, S. Stone, Prognostic value of an RNA expression signature derived from cell cycle proliferation genes in patients with prostate cancer: a retrospective study. The Lancet Oncology 12, 245–255 (2011).

4. E. A. Klein, Z. Haddad, K. Yousefi, L. L. C. Lam, Q. Wang, V. Choeurng, B. Palmer-Aronsten, C. Buerki, E. Davicioni, J. Li, M. W. Kattan, A. J. Stephenson, C. Magi-Galluzzi, Decipher Genomic Classifier Measured on Prostate Biopsy Predicts Metastasis Risk. Urology 90, 148–152 (2016).

5. C. J. Ryan, M. R. Smith, K. Fizazi, F. Saad, P. F. A. Mulders, C. N. Sternberg, K. Miller, C. J. Logothetis, N. D. Shore, E. J. Small, J. Carles, T. W. Flaig, M.-E. Taplin, C. S. Higano, P. de Souza, J. S. de Bono, T. W. Griffin, P. De Porre, M. K. Yu, Y. C. Park, J. Li, T. Kheoh, V. Naini, A. Molina, D. E. Rathkopf, Abiraterone acetate plus prednisone versus placebo plus prednisone in chemotherapy-naive men with metastatic castration-resistant prostate cancer (COU-AA-302): final overall survival analysis of a randomised, double-blind, placebo-controlled phase 3 study. The Lancet Oncology 16, 152–160 (2015).

6. Y. Loriot, K. Miller, C. N. Sternberg, K. Fizazi, J. S. De Bono, S. Chowdhury, C. S. Higano, S. Noonberg, S. Holmstrom, H. Mansbach, F. G. Perabo, D. Phung, C. Ivanescu, K. Skaltsa, T. M. Beer, B. Tombal, Effect of enzalutamide on health-related quality of life, pain, and skeletal-related events in asymptomatic and minimally symptomatic, chemotherapy-naive patients with metastatic castration-resistant prostate cancer (PREVAIL): results from a randomised, phase 3 trial. The Lancet Oncology 16, 509–521 (2015).

7. J. Mateo, S. Carreira, S. Sandhu, S. Miranda, H. Mossop, R. Perez-Lopez, D. Nava Rodrigues, D. Robinson, A. Omlin, N. Tunariu, G. Boysen, N. Porta, P. Flohr, A. Gillman, I. Figueiredo, C. Paulding, G. Seed, S. Jain, C. Ralph, A. Protheroe, S. Hussain, R. Jones, T. Elliott, U. McGovern, D. Bianchini, J. Goodall, Z. Zafeiriou, T. Williamson Chris, R. Ferraldeschi, R. Riisnaes, B. Ebbs, G. Fowler, D. Roda, W. Yuan, Y.-M. Wu, X. Cao, R. Brough, H. Pemberton, R. A’Hern, A. Swain, P. Kunju Lakshmi, R. Eeles, G. Attard, J. Lord Christopher, A. Ashworth, A. Rubin Mark, E. Knudsen Karen, Y. Feng Felix, M. Chinnaiyan Arul, E. Hall, S. de Bono Johann, DNA-Repair Defects and Olaparib in Metastatic Prostate Cancer. New England Journal of Medicine 373, 1697–1708 (2015).

8. F. Karzai, D. VanderWeele, R. A. Madan, H. Owens, L. M. Cordes, A. Hankin, A. Couvillon, E. Nichols, M. Bilusic, M. L. Beshiri, K. Kelly, V. Krishnasamy, S. Lee, M.-J. Lee, A. Yuno, J. B. Trepel, M. J. Merino, R. Dittamore, J. Marté, R. N. Donahue, J. Schlom, K. J. Killian, P. S. Meltzer, S. M. Steinberg, J. L. Gulley, J.-M. Lee, W. L. Dahut, Activity of durvalumab plus olaparib in metastatic castration-resistant prostate cancer in men with and without DNA damage repair mutations. Journal for ImmunoTherapy of Cancer 6, 141–153 (2018).

9. E. Castro, C. Goh, D. Leongamornlert, E. Saunders, M. Tymrakiewicz, T. Dadaev, K. Govindasami, M. Guy, S. Ellis, D. Frost, E. Bancroft, T. Cole, M. Tischkowitz, M. J. Kennedy, J. Eason, C. Brewer, D. G. Evans, R. Davidson, D. Eccles, M. E. Porteous, F. Douglas, J. Adlard, A. Donaldson, A. C. Antoniou, Z. Kote-Jarai, D. F. Easton, D. Olmos, R. Eeles, Effect of BRCA Mutations on Metastatic Relapse and Cause-specific Survival After Radical Treatment for Localised Prostate Cancer. European Urology 68, 186–193 (2015).

10. H. Kaur, D. C. Salles, S. Murali, J. L. Hicks, M. Nguyen, C. C. Pritchard, A. M. De Marzo, J. S. Lanchbury, B. J. Trock, W. B. Isaacs, K. M. Timms, E. S. Antonarakis, T. L. Lotan, Genomic and Clinicopathologic Characterization of ATM-deficient Prostate Cancer. Clinical Cancer Research 26, 4869–4881 (2020).

11. J. Armenia, S. A. M. Wankowicz, D. Liu, J. Gao, R. Kundra, E. Reznik, W. K. Chatila, D. Chakravarty, G. C. Han, I. Coleman, B. Montgomery, C. Pritchard, C. Morrissey, C. E. Barbieri, H. Beltran, A. Sboner, Z. Zafeiriou, S. Miranda, C. M. Bielski, A. V. Penson, C. Tolonen, F. W. Huang, D. Robinson, Y. M. Wu, R. Lonigro, L. A. Garraway, F. Demichelis, P. W. Kantoff, M.-E. Taplin, W. Abida, B. S. Taylor, H. I. Scher, P. S. Nelson, J. S. de Bono, M. A. Rubin, C. L. Sawyers, A. M. Chinnaiyan, N. Schultz, E. M. Van Allen, P. S. C. I. P. C. D. Team, The long tail of oncogenic drivers in prostate cancer. Nature Genetics 50, 645–651 (2018).

12. C. G. A. R. Network, The Molecular Taxonomy of Primary Prostate Cancer. Cell 163, 1011–1025 (2015).

13. C. S. Cooper, R. Eeles, D. C. Wedge, P. Van Loo, G. Gundem, L. B. Alexandrov, B. Kremeyer, A. Butler, A. G. Lynch, N. Camacho, C. E. Massie, J. Kay, H. J. Luxton, S. Edwards, Z. Kote-Jarai, N. Dennis, S. Merson, D. Leongamornlert, J. Zamora, C. Corbishley, S. Thomas, S. Nik-Zainal, M. Ramakrishna, S. O’Meara, L. Matthews, J. Clark, R. Hurst, R. Mithen, R. G. Bristow, P. C. Boutros, M. Fraser, S. Cooke, K. Raine, D. Jones, A. Menzies, L. Stebbings, J. Hinton, J. Teague, S. McLaren, L. Mudie, C. Hardy, E. Anderson, O. Joseph, V. Goody, B. Robinson, M. Maddison, S. Gamble, C. Greenman, D. Berney, S. Hazell, N. Livni, C. Fisher, C. Ogden, P. Kumar, A. Thompson, C. Woodhouse, D. Nicol, E. Mayer, T. Dudderidge, N. C. Shah, V. Gnanapragasam, T. Voet, P. Campbell, A. Futreal, D. Easton, A. Y. Warren, C. S. Foster, M. R. Stratton, H. C. Whitaker, U. McDermott, D. S. Brewer, D. E. Neal, I. P. G. the, Analysis of the genetic phylogeny of multifocal prostate cancer identifies multiple independent clonal expansions in neoplastic and morphologically normal prostate tissue. Nature Genetics 47, 367–372 (2015).

14. M. Fraser, V. Y. Sabelnykova, T. N. Yamaguchi, L. E. Heisler, J. Livingstone, V. Huang, Y.-J. Shiah, F. Yousif, X. Lin, A. P. Masella, N. S. Fox, M. Xie, S. D. Prokopec, A. Berlin, E. Lalonde, M. Ahmed, D. Trudel, X. Luo, T. A. Beck, A. Meng, J. Zhang, A. D’Costa, R. E. Denroche, H. Kong, S. M. G. Espiritu, M. L. K. Chua, A. Wong, T. Chong, M. Sam, J. Johns, L. Timms, N. B. Buchner, M. Orain, V. Picard, H. Hovington, A. Murison, K. Kron, N. J. Harding, C. P’ng, K. E. Houlahan, K. C. Chu, B. Lo, F. Nguyen, C. H. Li, R. X. Sun, R. de Borja, C. I. Cooper, J. F. Hopkins, S. K. Govind, C. Fung, D. Waggott, J. Green, S. Haider, M. A. Chan-Seng-Yue, E. Jung, Z. Wang, A. Bergeron, A. Dal Pra, L. Lacombe, C. C. Collins, C. Sahinalp, M. Lupien, N. E. Fleshner, H. H. He, Y. Fradet, B. Tetu, T. van der Kwast, J. D. McPherson, R. G. Bristow, P. C. Boutros, Genomic hallmarks of localized, non-indolent prostate cancer. Nature 541, 359–364 (2017).

15. G. Gundem, P. Van Loo, B. Kremeyer, L. B. Alexandrov, J. M. C. Tubio, E. Papaemmanuil, D. S. Brewer, H. M. L. Kallio, G. Högnäs, M. Annala, K. Kivinummi, V. Goody, C. Latimer, S. O’Meara, K. J. Dawson, W. Isaacs, M. R. Emmert-Buck, M. Nykter, C. Foster, Z. Kote-Jarai, D. Easton, H. C. Whitaker, D. E. Neal, C. S. Cooper, R. A. Eeles, T. Visakorpi, P. J. Campbell, U. McDermott, D. C. Wedge, G. S. Bova, I. P. U. Group, The evolutionary history of lethal metastatic prostate cancer. Nature 520, 353–357 (2015).

16. C. Gerhauser, F. Favero, T. Risch, R. Simon, L. Feuerbach, Y. Assenov, D. Heckmann, N. Sidiropoulos, S. M. Waszak, D. Hübschmann, A. Urbanucci, E. G. Girma, V. Kuryshev, L. J. Klimczak, N. Saini, A. M. Stütz, D. Weichenhan, L.-M. Böttcher, R. Toth, J. D. Hendriksen, C. Koop, P. Lutsik, S. Matzk, H.-J. Warnatz, V. Amstislavskiy, C. Feuerstein, B. Raeder, O. Bogatyrova, E.-M. Schmitz, C. Hube-Magg, M. Kluth, H. Huland, M. Graefen, C. Lawerenz, G. H. Henry, T. N. Yamaguchi, A. Malewska, J. Meiners, D. Schilling, E. Reisinger, R. Eils, M. Schlesner, D. W. Strand, R. G. Bristow, P. C. Boutros, C. von Kalle, D. Gordenin, H. Sültmann, B. Brors, G. Sauter, C. Plass, M.-L. Yaspo, J. O. Korbel, T. Schlomm, J. Weischenfeldt, Molecular Evolution of Early-Onset Prostate Cancer Identifies Molecular Risk Markers and Clinical Trajectories. Cancer Cell 34, 996–1011 (2018).

17. N. C. I. Surveillance Research Program. (2024), chap. 2024 Apr 17.

18. M. T. Chang, S. Asthana, S. P. Gao, B. H. Lee, J. S. Chapman, C. Kandoth, J. Gao, N. D. Socci, D. B. Solit, A. B. Olshen, N. Schultz, B. S. Taylor, Identifying recurrent mutations in cancer reveals widespread lineage diversity and mutational specificity. Nature Biotechnology 34, 155–163 (2016).

19. C. Mitsopoulos, P. Di Micco, E. V. Fernandez, D. Dolciami, E. Holt, I. L. Mica, E. A. Coker, J. E. Tym, J. Campbell, K. H. Che, B. Ozer, C. Kannas, A. A. Antolin, P. Workman, B. Al-Lazikani, canSAR: update to the cancer translational research and drug discovery knowledgebase. Nucleic Acids Research 49, 1074–1082 (2020).

20. D. C. Wedge, G. Gundem, T. Mitchell, D. J. Woodcock, I. Martincorena, M. Ghori, J. Zamora, A. Butler, H. Whitaker, Z. Kote-Jarai, L. B. Alexandrov, P. Van Loo, C. E. Massie, S. Dentro, A. Y. Warren, C. Verrill, D. M. Berney, N. Dennis, S. Merson, …, T. C. The, Sequencing of prostate cancers identifies new cancer genes, routes of progression and drug targets. Nature Genetics 50, 682–692 (2018).

21. P. di Micco, A. A. Antolin, C. Mitsopoulos, E. Villasclaras-Fernandez, D. Sanfelice, D. Dolciami, P. Ramagiri, Ioan L. Mica, Joseph E. Tym, Philip W. Gingrich, H. Hu, P. Workman, B. Al-Lazikani, canSAR: update to the cancer translational research and drug discovery knowledgebase. Nucleic Acids Research 51, 1212–1219 (2023).

22. D. Burns, E. Anokian, E. J. Saunders, R. G. Bristow, M. Fraser, J. Reimand, T. Schlomm, G. Sauter, B. Brors, J. Korbel, J. Weischenfeldt, S. M. Waszak, N. M. Corcoran, C.-H. Jung, B. J. Pope, C. M. Hovens, G. Cancel-Tassin, O. Cussenot, M. Loda, …, R. A. Eeles, Rare Germline Variants Are Associated with Rapid Biochemical Recurrence After Radical Prostate Cancer Treatment: A Pan Prostate Cancer Group Study. European Urology 82, 201–211 (2022).

23. J. M. Dempster, I. Boyle, F. Vazquez, D. E. Root, J. S. Boehm, W. C. Hahn, A. Tsherniak, J. M. McFarland, Chronos: a cell population dynamics model of CRISPR experiments that improves inference of gene fitness effects. Genome Biology 22, 343–366 (2021).

24. W. Yang, J. Soares, P. Greninger, E. J. Edelman, H. Lightfoot, S. Forbes, N. Bindal, D. Beare, J. A. Smith, I. R. Thompson, S. Ramaswamy, P. A. Futreal, D. A. Haber, M. R. Stratton, C. Benes, U. McDermott, M. J. Garnett, Genomics of Drug Sensitivity in Cancer (GDSC): a resource for therapeutic biomarker discovery in cancer cells. Nucleic Acids Research 41, 955–961 (2013).

25. R. T. K. Poluri, C. J. Beauparlant, A. Droit, É. Audet-Walsh, RNA sequencing data of human prostate cancer cells treated with androgens. Data in Brief 25, 1–4 (2019).

26. J. Cham, A. R. Venkateswaran, M. Bhangoo, Targeting the PI3K-AKT-mTOR Pathway in Castration Resistant Prostate Cancer: A Review Article. Clinical Genitourinary Cancer 19, 563.e561–563.e567 (2021).

27. B. Y. Shorning, M. S. Dass, M. J. Smalley, H. B. Pearson, The PI3K-AKT-mTOR Pathway and Prostate Cancer: At the Crossroads of AR, MAPK, and WNT Signaling. International Journal of Molecular Sciences. 2020 (10.3390/ijms21124507).

28. D. Basak, L. Gregori, F. Johora, S. Deb, Preclinical and Clinical Research Models of Prostate Cancer: A Brief Overview. Life. 2022 (10.3390/life12101607).

29. S. L. Wang, Z. Liao, A. A. Vaporciyan, S. L. Tucker, H. Liu, X. Wei, S. Swisher, J. A. Ajani, J. D. Cox, R. Komaki, Investigation of clinical and dosimetric factors associated with postoperative pulmonary complications in esophageal cancer patients treated with concurrent chemoradiotherapy followed by surgery. Int J Radiat Oncol Biol Phys 64, 692–699 (2006).

30. Z. Fan, B. Yin, X. Chen, G. Yang, W. Zhang, X. Ye, H. Han, M. Li, M. Shu, R. Liu, Comprehensive analysis of paraspeckle-associated gene modules unveils prognostic signatures and immunological relevance in multi-cancers. Discover Oncology 15, 345–362 (2024).

31. W.-q. Du, Z.-m. Zhu, X. Jiang, M.-j. Kang, D.-s. Pei, COPS6 promotes tumor progression and reduces CD8+ T cell infiltration by repressing IL-6 production to facilitate tumor immune evasion in breast cancer. Acta Pharmacologica Sinica 44, 1890–1905 (2023).

32. X. Fan, X. Yang, N. Guo, X. Gao, Y. Zhao, Development of an endoplasmic reticulum stress-related signature with potential implications in prognosis and immunotherapy in head and neck squamous cell carcinoma. Diagnostic Pathology 18, 1–14 (2023).

33. H.-Q. Yu, F. Li, H. Xiong, L. Fang, J. Zhang, P. Bie, C.-M. Xie, Elevated FBXL18 promotes RPS15A ubiquitination and SMAD3 activation to drive HCC. Hepatology Communications 7, 1–15 (2023).

34. R. Tian, J. Tian, X. Zuo, S. Ren, H. Zhang, H. Liu, Z. Wang, Y. Cui, R. Niu, F. Zhang, RACK1 facilitates breast cancer progression by competitively inhibiting the binding of β-catenin to PSMD2 and enhancing the stability of β-catenin. Cell Death & Disease 14, 1–13 (2023).

35. J. M. Dolezal, A. P. Dash, E. V. Prochownik, Diagnostic and prognostic implications of ribosomal protein transcript expression patterns in human cancers. BMC Cancer 18, 1–14 (2018).

36. W. McLaren, L. Gil, S. E. Hunt, H. S. Riat, G. R. S. Ritchie, A. Thormann, P. Flicek, F. Cunningham, The Ensembl Variant Effect Predictor. Genome Biology 17, 122–136 (2016).

37. J. S. Myers, A. K. von Lersner, C. J. Robbins, Q.-X. A. Sang, Differentially Expressed Genes and Signature Pathways of Human Prostate Cancer. Public Library of Science One 10, 1–27 (2015).

38 D. Broad, DepMap 22Q4 Public. (2022).

