## Supplemental Materials for "Distinct druggable biological processes in early-onset prostate cancer"

**Supplementary Content**

### Supplementary Figures

#### Figure S1: Expanded study workflow

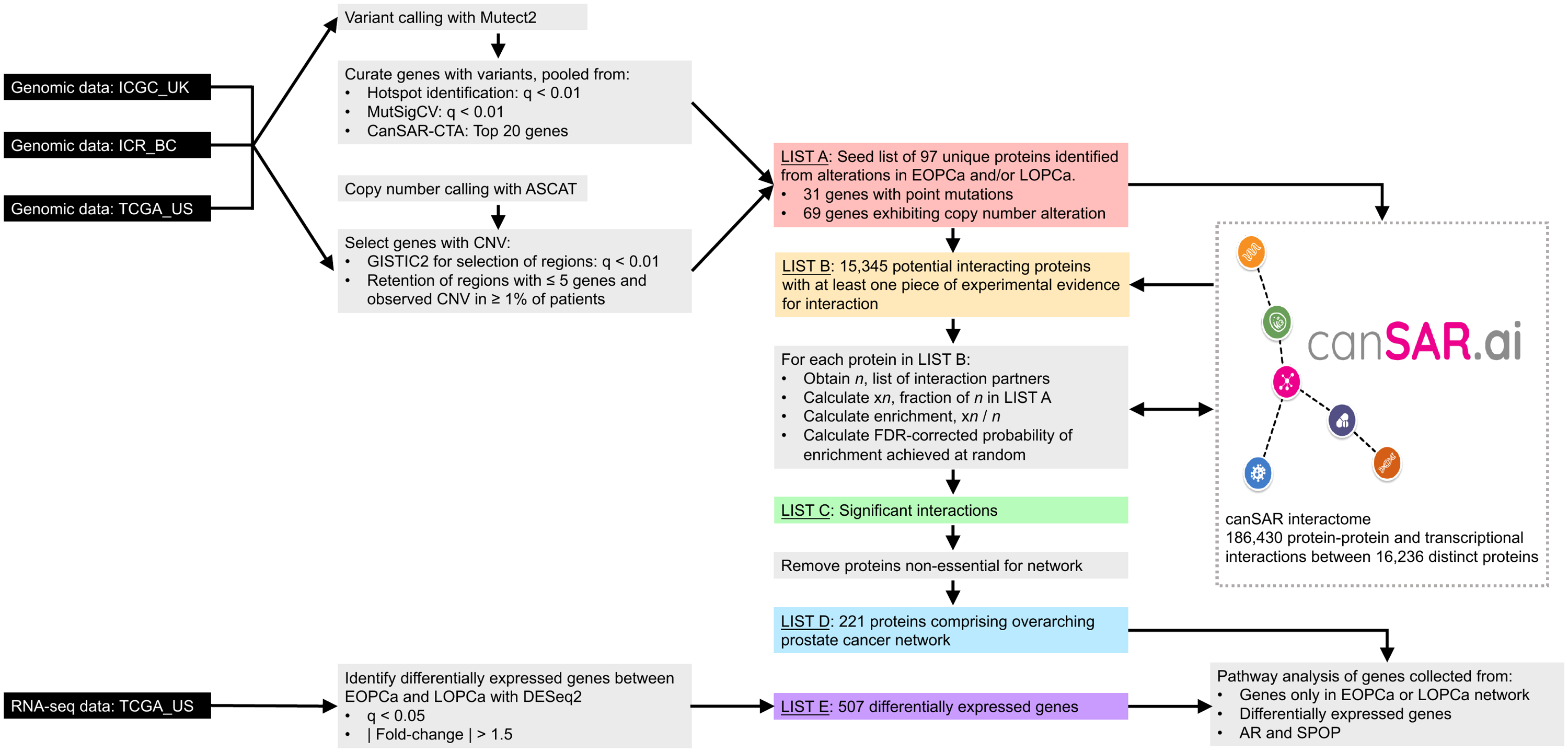

#### Figure S2: Network of differential WES and RNA-seq genes annotated by pathway cluster: EOPCa

Colors used follow those in Figure 2.

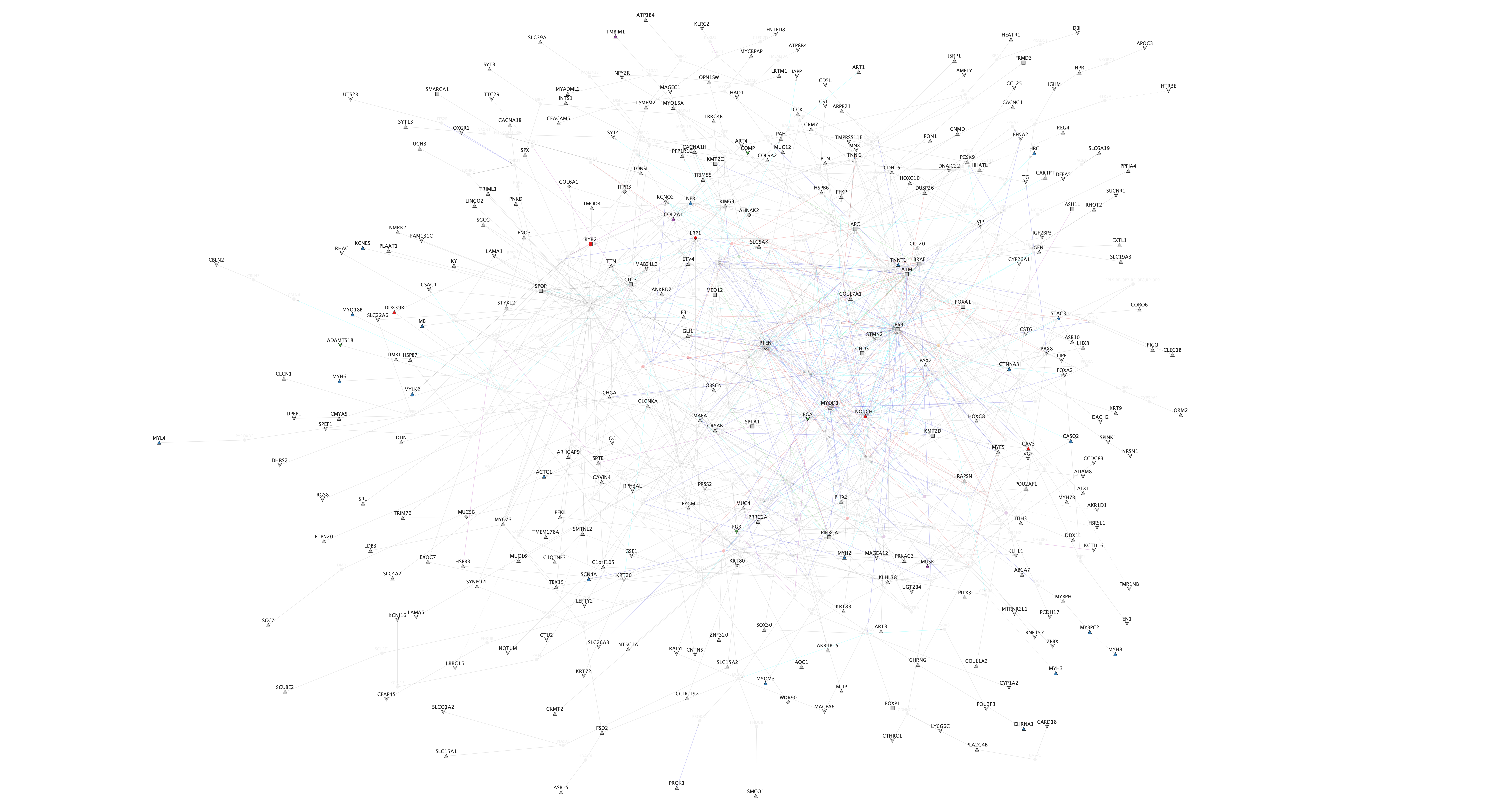

#### Figure S3: Network of differential WES and RNA-seq genes annotated by pathway cluster: LOPCa

Colors used follow those in Figure 2.

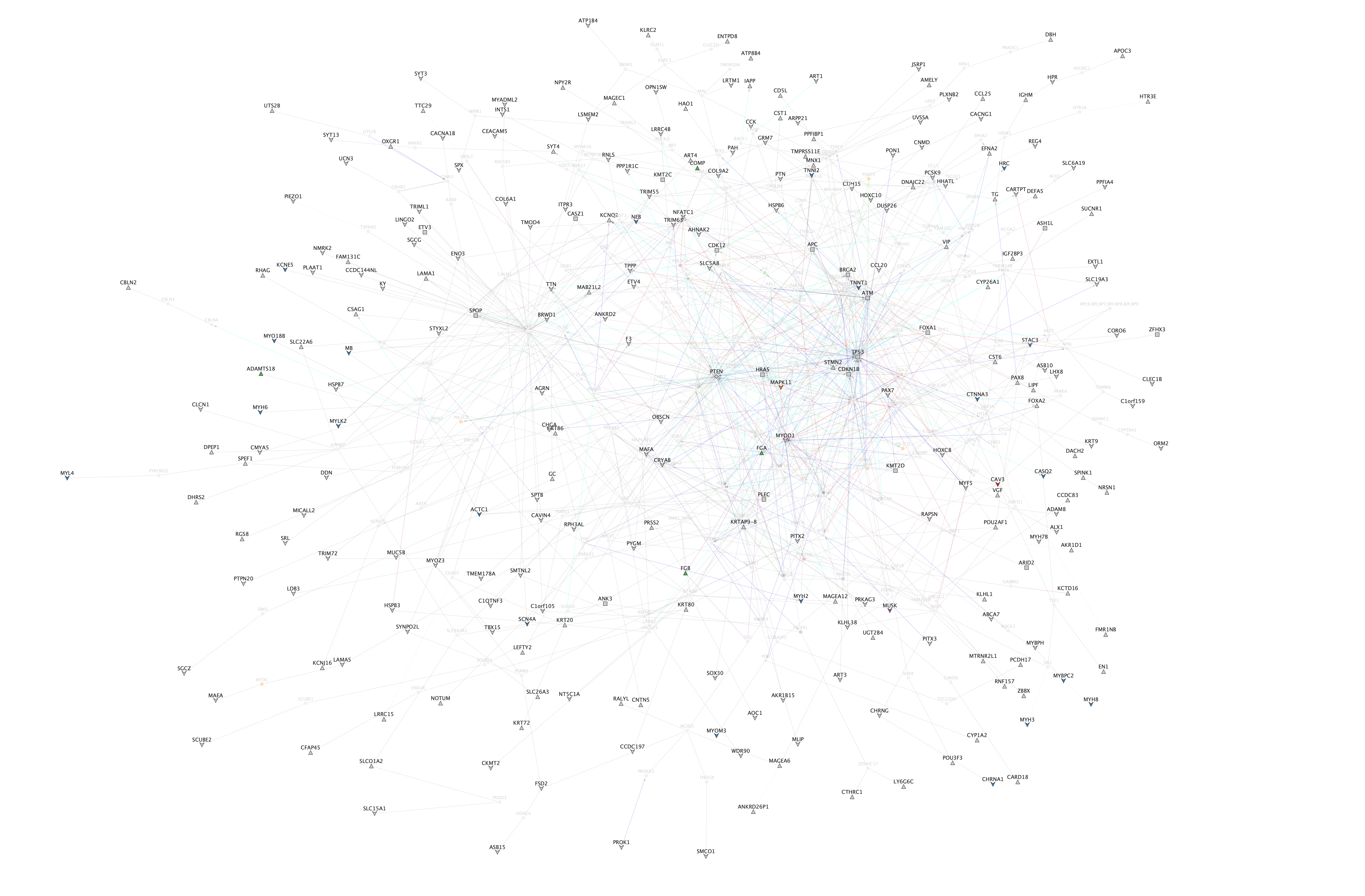

#### Figure S4: Network of differential WES and RNA-seq genes annotated by druggability: EOPCa

Colors used follow those in Figure 2.

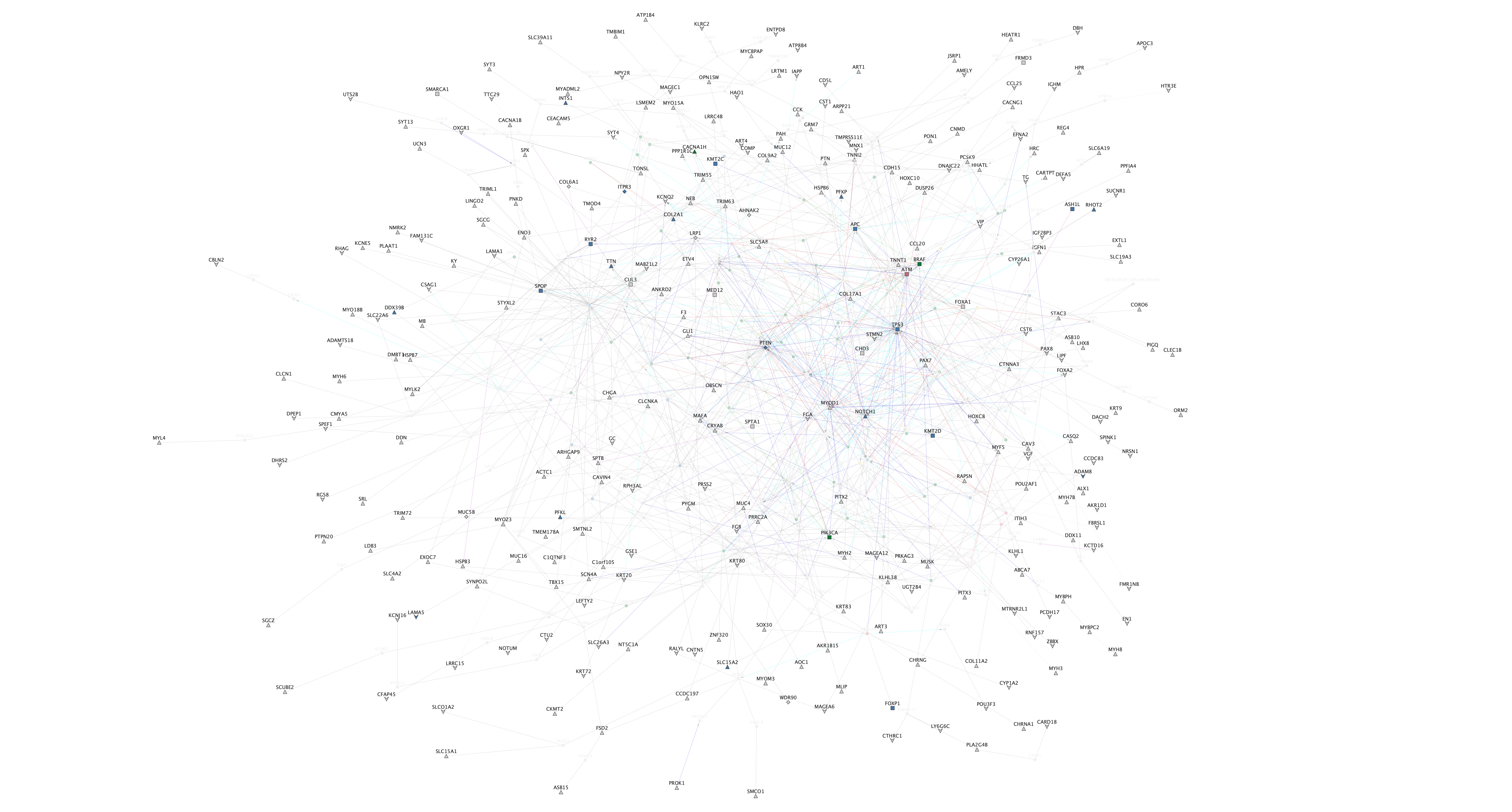

#### Figure S5: Network of differential WES and RNA-seq genes annotated by druggability: LOPCa

Colors used follow those in Figure 2.

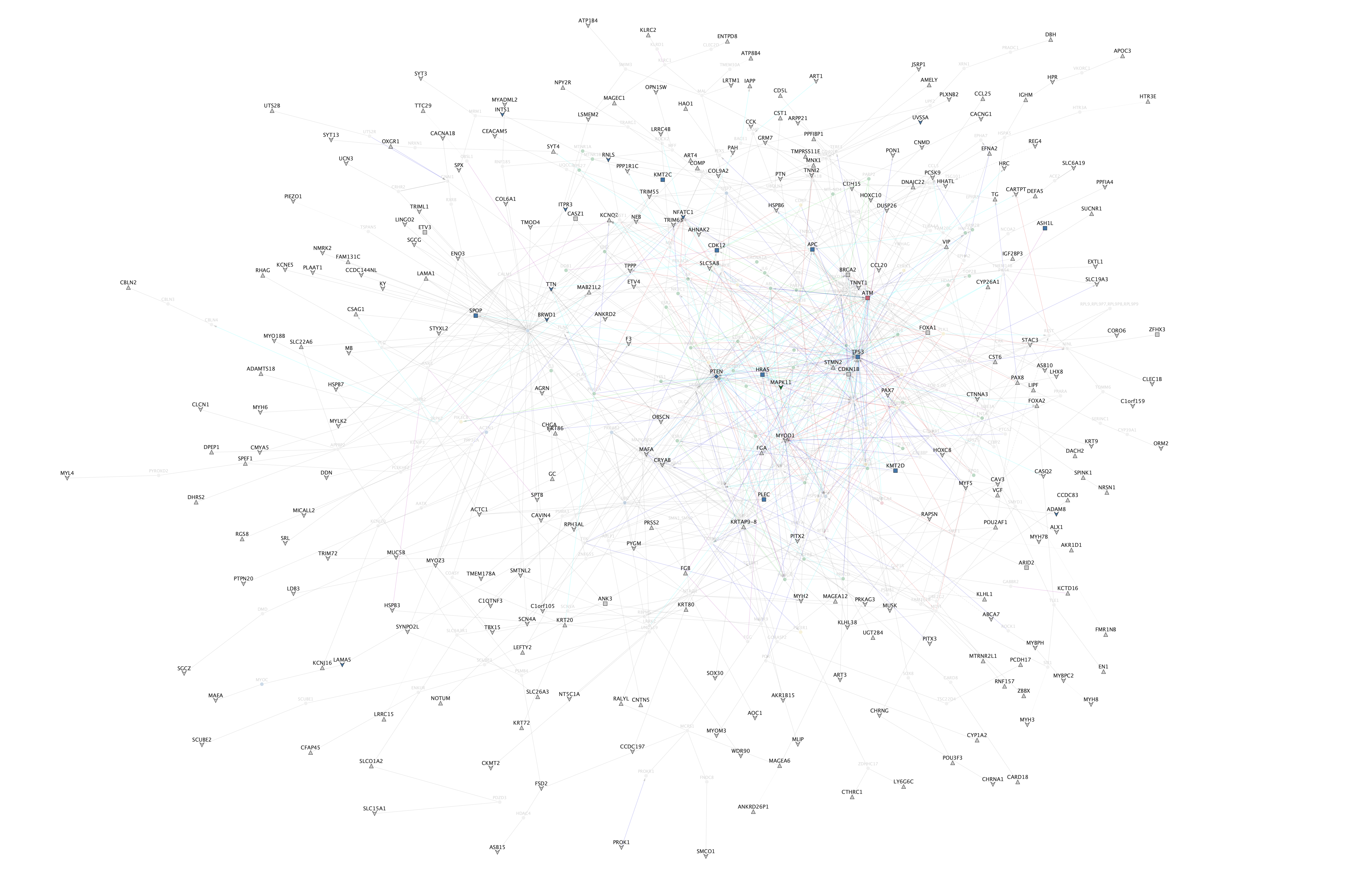

#### Figure S6: Pathway annotation of differential WES network genes and differentially expressed genes

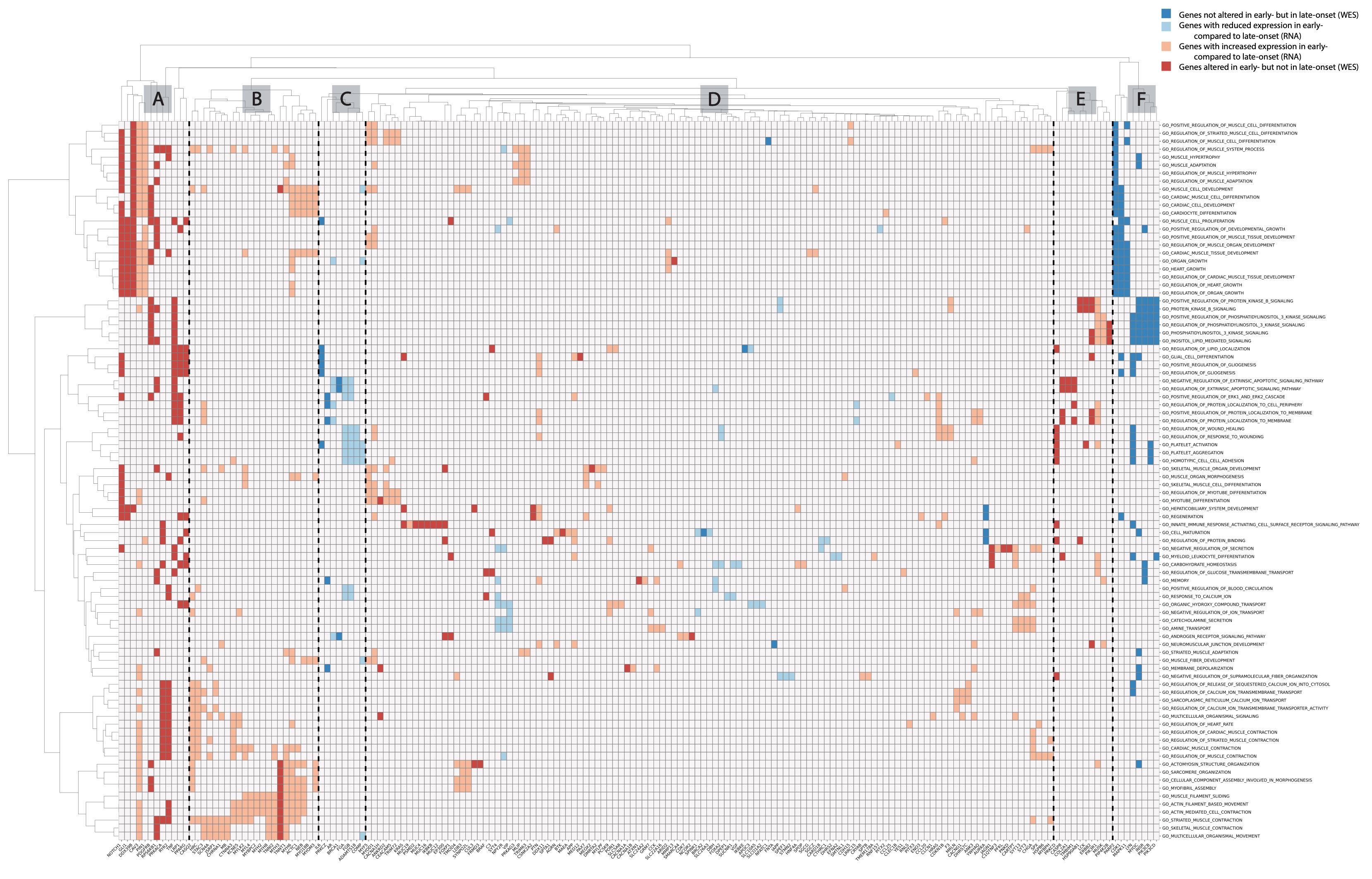

#### Figure S7: Number of genetic dependencies identified using a gene set of the same size of the one in this study (randomly selected 221 genes).

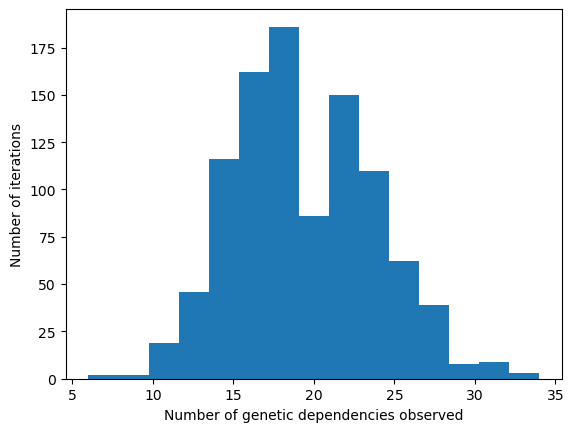

#### Figure S8: Venn diagram of genes identified from WES alterations, identified by differential expression, and included in the main networks

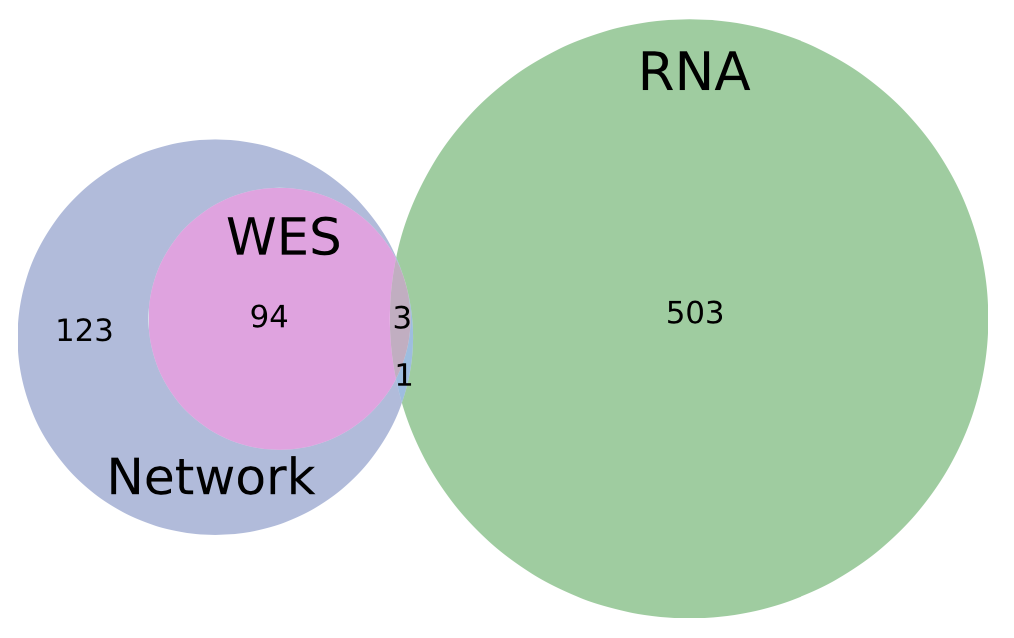

### Supplementary Notes

#### Supplementary Note 1: Significance of the number of dependencies identified

To verify that this number of dependencies identified in our study is significant compared to a random gene set of the same size, we performed the following statistical experiment. Using the full human genome (17,386 genes), we generated 1000 lists of 221 randomly selected genes. For these, we tested how many of these random proteins were dependencies in PCa cell lines using the same criteria we used for the true set of 221 derived from our analysis. **Fig. S7** shows the distribution of the number of dependencies observed for each of these 1000 random trials. The range is 5-35 and the mode is 17 dependencies. Our ‘true’ observed genetic dependencies of 39 are not observed.

### Supplementary Tables

#### Table S1: Interventional clinical trials from clinicaltrials.gov for prostate cancer with patients harboring PTEN loss

| NCT Number | Study Title | Study Status | Interventions | Phases |
| --- | --- | --- | --- | --- |
| NCT00071968 | Neoadjuvant CCI-779 Followed By Radical Prostatectomy in Treating Patients With Newly Diagnosed Prostate Cancer Who Have a High Risk of Relapse | COMPLETED | DRUG: temsirolimus\|PROCEDURE: conventional surgery\|PROCEDURE: neoadjuvant therapy | PHASE2 |
| NCT00078910 | Neoadjuvant Exisulind in Treating Patients Who Are Undergoing Radical Prostatectomy for Stage II or Stage III Prostate Cancer | COMPLETED | DRUG: exisulind\|PROCEDURE: conventional surgery\|PROCEDURE: neoadjuvant therapy | PHASE2 |
| NCT00235794 | An Open Label Exploratory Study in Newly Diagnosed Prostate Cancer Patients | COMPLETED | DRUG: temsirolimus | NA |
| NCT00311623 | Sirolimus Before Surgery in Treating Patients With Advanced Localized Prostate Cancer | COMPLETED | DRUG: Rapamycin 3mg\|DRUG: Rapamycin 6mg\|PROCEDURE: Radical prostatectomy | PHASE1\|PHASE2 |
| NCT00579514 | Germline Alterations of Tumor Susceptibility Genes in New York Cancer Patients | RECRUITING | GENETIC: PCR/PCR/LDR Strategy | NA |
| NCT00657982 | Phase II Study of RAD001 in a Neoadjuvant Setting in Men With Intermediate or High Risk Prostate Cancer | UNKNOWN | DRUG: RAD001 | PHASE2 |
| NCT00881725 | A Study of Pre-operative Metformin in Prostate Cancer | TERMINATED | DRUG: Metformin | PHASE2 |
| NCT01009736 | Effects of Tomato-Soy Juice on Biomarkers in Patients With Prostate Cancer Undergoing Prostatectomy | COMPLETED | DIETARY_SUPPLEMENT: tomato-soy juice\|OTHER: laboratory biomarker analysis\|OTHER: pharmacological study\|PROCEDURE: therapeutic conventional surgery | PHASE1\|PHASE2 |
| NCT01075308 | SB939 in Treating Patients With Recurrent or Metastatic Prostate Cancer | COMPLETED | DRUG: HDAC inhibitor SB939 | PHASE2 |
| NCT01444820 | Hypofractionated, Dose Escalation Radiotherapy for High Risk Adenocarcinoma of the Prostate | UNKNOWN | RADIATION: hypofractionation\|RADIATION: conventional | PHASE3 |
| NCT01485861 | Study of Ipatasertib or Apitolisib With Abiraterone Acetate Versus Abiraterone Acetate in Participants With Castration-Resistant Prostate Cancer Previously Treated With Docetaxel Chemotherapy | COMPLETED | DRUG: Abiraterone\|DRUG: Apitolisib\|DRUG: Ipatasertib\|DRUG: Placebo\|DRUG: Prednisone\|DRUG: Prednisolone | PHASE1\|PHASE2 |
| NCT01542021 | Androgen Deprivation Therapy Prior to Prostatectomy for Patients With Intermediate and High Risk Prostate Cancer | ACTIVE_NOT_RECRUITING | DRUG: degarelix injection\|DRUG: degarelix injection\|DRUG: androgen deprivation therapy | NA |
| NCT01681433 | OGX-427 in Metastatic Castrate-Resistant Prostate Cancer With Prostate-Specific Antigen Progression While Receiving Abiraterone | TERMINATED | DRUG: OGX-427\|DRUG: Abiraterone Acetate\|DRUG: Prednisone | PHASE2 |
| NCT01717898 | A Multicenter Phase I/II Trial of Abiraterone Acetate + BEZ235 in Metastatic, Castration-Resistant Prostate Cancer | TERMINATED | DRUG: BEZ235\|DRUG: Prednisone\|DRUG: Abiraterone acetate | PHASE1\|PHASE2 |
| NCT01741753 | BKM120+Abiraterone Acetate for Metastatic CRPC | TERMINATED | DRUG: BKM120\|DRUG: Abiraterone\|DRUG: Prednisone | PHASE1 |
| NCT01751451 | 3-arm Study of Abiraterone Acetate Alone, Abiraterone Acetate Plus Degarelix, a GnRH Antagonist, and Degarelix Alone for Patients With Prostate Cancer With a Rising PSA or a Rising PSA and Nodal Disease Following Definitive Radical Prostatectomy | COMPLETED | DRUG: Abiraterone acetate\|DRUG: Abiraterone acetate plus degarelix\|DRUG: Degarelix | PHASE2 |
| NCT01884285 | AZD8186 First Time In Patient Ascending Dose Study | COMPLETED | DRUG: Part A: AZD8186 monotherapy\|DRUG: Part B: AZD8186 monotherapy\|DRUG: Part C1: Abiraterone acetate combination with AZD8186\|DRUG: Part D1: AZD2014 combination with AZD8186\|DRUG: Part D2 AZD2014 combination with AZD8186\|DRUG: Part C2: Abiraterone acetate combination with AZD8186 | PHASE1 |
| NCT02090114 | RE-sensitizing With Supraphysiologic Testosterone to Overcome REsistance (The RESTORE Study) | COMPLETED | DRUG: Testosterone cypionate\|DRUG: Testosterone Enanthate\|DRUG: Abiraterone acetate\|DRUG: Enzalutamide | PHASE2 |
| NCT02099864 | Genetic and Molecular Mechanisms in Assessing Response in Patients With Prostate Cancer Receiving Enzalutamide Therapy | ACTIVE_NOT_RECRUITING | DRUG: Enzalutamide | PHASE2 |
| NCT02159690 | A Phase II Neoadjuvant Study of Enzalutamide, Abiraterone Acetate, Dutasteride and Degarelix in Men With Localized Prostate Cancer Pre-prostatectomy | WITHDRAWN | DRUG: Enzalutamide\|DRUG: Abiraterone acetate\|DRUG: Prednisone\|DRUG: Dutasteride\|DRUG: Degarelix | PHASE2 |
| NCT02215096 | Dose-finding Study of GSK2636771 When Administered in Combination With Enzalutamide in Male Subjects With Metastatic Castration-Resistant Prostate Cancer | COMPLETED | DRUG: GSK2636771\|DRUG: Enzalutamide | PHASE1 |
| NCT02303327 | Comparative Study of Radiotherapy Treatments to Treat High Risk Prostate Cancer Patients | RECRUITING | RADIATION: EBRT + HDR brachytherapy boost\|RADIATION: Hypofractionated Dose Escalation Radiotherapy\|DRUG: Androgen deprivation therapy | PHASE3 |
| NCT02311764 | Carboplatin in Castration-resistant Prostate Cancer | TERMINATED | DRUG: Carboplatin | PHASE2 |
| NCT02430480 | Using Multiparametric MRI to Evaluate Intraprostatic Tumor Responses and Androgen Resistance Patterns in Newly Diagnosed Prostate Cancer | ACTIVE_NOT_RECRUITING | DRUG: Goserelin\|DRUG: Enzalutamide\|DEVICE: mpMRI | PHASE2 |
| NCT02465060 | Targeted Therapy Directed by Genetic Testing in Treating Patients With Advanced Refractory Solid Tumors, Lymphomas, or Multiple Myeloma (The MATCH Screening Trial) | ACTIVE_NOT_RECRUITING | DRUG: Adavosertib\|DRUG: Afatinib\|DRUG: Afatinib Dimaleate\|DRUG: Binimetinib\|PROCEDURE: Biopsy\|PROCEDURE: Biospecimen Collection\|DRUG: Capivasertib\|PROCEDURE: Computed Tomography\|DRUG: Copanlisib\|DRUG: Copanlisib Hydrochloride\|DRUG: Crizotinib\|OTHER: Cytology Specimen Collection Procedure\|DRUG: Dabrafenib\|DRUG: Dabrafenib Mesylate\|DRUG: Dasatinib\|DRUG: Defactinib\|DRUG: Defactinib Hydrochloride\|PROCEDURE: Echocardiography\|DRUG: Erdafitinib\|DRUG: Fexagratinib\|DRUG: Ipatasertib\|OTHER: Laboratory Biomarker Analysis\|DRUG: Larotrectinib\|DRUG: Larotrectinib Sulfate\|PROCEDURE: Magnetic Resonance Imaging\|BIOLOGICAL: Nivolumab\|DRUG: Osimertinib\|DRUG: Palbociclib\|BIOLOGICAL: Pertuzumab\|DRUG: PI3K-beta Inhibitor GSK2636771\|PROCEDURE: Radiologic Examination\|BIOLOGICAL: Relatlimab\|DRUG: Sapanisertib\|DRUG: Sunitinib Malate\|DRUG: Taselisib\|DRUG: Trametinib\|BIOLOGICAL: Trastuzumab\|BIOLOGICAL: Trastuzumab Emtansine\|DRUG: Ulixertinib\|DRUG: Vismodegib | PHASE2 |
| NCT02478125 | Pilot Study of Mobilization and Treatment of Disseminated Tumor Cells in Men With Metastatic Prostate Cancer | TERMINATED | DRUG: Burixafor Hydrobromide\|DRUG: Docetaxel\|DRUG: G-CSF | PHASE1 |
| NCT02821728 | Sulphate Accumulation in Prostate | COMPLETED | DIETARY_SUPPLEMENT: Dietary intervention\|OTHER: Normal diet | NA |
| NCT02961257 | Trial Evaluating the Safety of 2 Schedules of Cabazitaxel in Elderly Men With mCRPC Previously Treated With a Docetaxel | COMPLETED | DRUG: cabazitaxel\|DRUG: Prednisone\|DRUG: Granulocyte colony-stimulating factor (G-CSF) | PHASE3 |
| NCT03072238 | Ipatasertib Plus Abiraterone Plus Prednisone/Prednisolone, Relative to Placebo Plus Abiraterone Plus Prednisone/Prednisolone in Adult Male Patients With Metastatic Castrate-Resistant Prostate Cancer | ACTIVE_NOT_RECRUITING | DRUG: Ipatasertib\|DRUG: Abiraterone\|DRUG: Placebo\|DRUG: Prednisone/Prednisolone | PHASE3 |
| NCT03080116 | Neoadjuvant Degarelix With or Without Apalutamide (ARN-509) Followed by Radical Prostatectomy | UNKNOWN | DRUG: ARN-509\|DRUG: Degarelix\|OTHER: Placebo | PHASE2 |
| NCT03218826 | PI3Kbeta Inhibitor AZD8186 and Docetaxel in Treating Patients Advanced Solid Tumors With PTEN or PIK3CB Mutations That Are Metastatic or Cannot Be Removed by Surgery | ACTIVE_NOT_RECRUITING | DRUG: Docetaxel\|OTHER: Laboratory Biomarker Analysis\|OTHER: Pharmacological Study\|DRUG: PI3Kbeta Inhibitor AZD8186 | PHASE1 |
| NCT03572478 | Rucaparib and Nivolumab in Patients With Prostate or Endometrial Cancer | TERMINATED | DRUG: Rucaparib\|DRUG: Nivolumab | PHASE1\|PHASE2 |
| NCT03665922 | Biomarkers of Sulforaphane/Broccoli Sprout Extract in Prostate Cancer | ACTIVE_NOT_RECRUITING | DIETARY_SUPPLEMENT: BroccoMax¬Æ\|OTHER: Placebo | NA |
| NCT03673787 | A Trial of Ipatasertib in Combination With Atezolizumab | UNKNOWN | DRUG: ipatasertib\|DRUG: Atezolizumab | PHASE1\|PHASE2 |
| NCT04060394 | Dose-Escalation and Efficacy Study of LAE001/Prednisone Plus Afuresertib Patients With m-CRPC | ACTIVE_NOT_RECRUITING | DRUG: Phase I and Phase II: LAE001/prednisone + afuresertib | PHASE1\|PHASE2 |
| NCT04458311 | Abiraterone Acetate in Combination With Tildrakizumab | ACTIVE_NOT_RECRUITING | DRUG: Abiraterone Acetate\|DRUG: Tildrakizumab | PHASE1\|PHASE2 |
| NCT04493853 | Capivasertib+Abiraterone as Treatment for Patients With Metastatic Hormone-sensitive Prostate Cancer and PTEN Deficiency | ACTIVE_NOT_RECRUITING | DRUG: Capivasertib\|OTHER: Placebo\|DRUG: Abiraterone Acetate | PHASE3 |
| NCT04586270 | A Study of TAS0612 in Participants With Advanced or Metastatic Solid Tumor Cancer | RECRUITING | DRUG: TAS0612 | PHASE1 |
| NCT04601441 | Study to Evaluate ctDNA of mCSPC Patients Receiving Apalutamide in Japan | RECRUITING | DRUG: Apalutamide | PHASE4 |
| NCT04737109 | Neoadjuvant Androgen Deprivation, Darolutamide, and Ipatasertib in Men With Localized, High Risk Prostate Cancer | TERMINATED | DRUG: Ipatasertib\|DRUG: Darolutamide\|DRUG: Androgen Deprivation Therapy | PHASE1\|PHASE2 |
| NCT05563558 | Pembrolizumab, Carboplatin and Cabazitaxel in Aggressive Metastatic Castration Resistant Prostate Cancer (PEAPOD_FOS) | RECRUITING | DRUG: Pembrolizumab\|DRUG: Carboplatin\|DRUG: Cabazitaxel | PHASE2 |
| NCT05593497 | A Single-Arm Phase II Study of Neoadjuvant Intensified Androgen Deprivation (Leuprolide and Abiraterone Acetate) in Combination With AKT Inhibition (Capivasertib) for High-Risk Localized Prostate Cancer With PTEN Loss | NOT_YET_RECRUITING | DRUG: Capivasertib\|DRUG: abiraterone acetate | PHASE2 |
| NCT06029998 | Bortezomib in Patients With Metastatic Castration-Resistant Prostate Cancer With PTEN Deletion | NOT_YET_RECRUITING | DRUG: Bortezomib | PHASE2 |
| NCT06183736 | CVL237 Tablets in the Treatment of Advanced Solid Tumors With PTEN Deficiency | NOT_YET_RECRUITING | DRUG: CVL237 tablets | PHASE2 |

#### Table S2: Description of 9 prostate cancer cell lines with CRISPR profiling data available

Age entries are left empty if information was not available; rows highlighted denote cell lines derived from patients <60 years old. Lines shown reflect those in 22Q4 DepMap release.

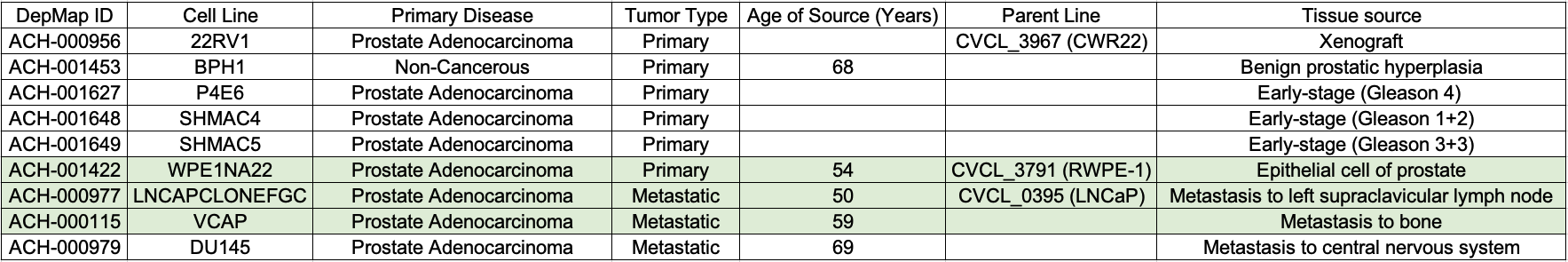

#### Table S3: Dependencies occurring in EOPCa observed in cell lines derived from EOPCa samples

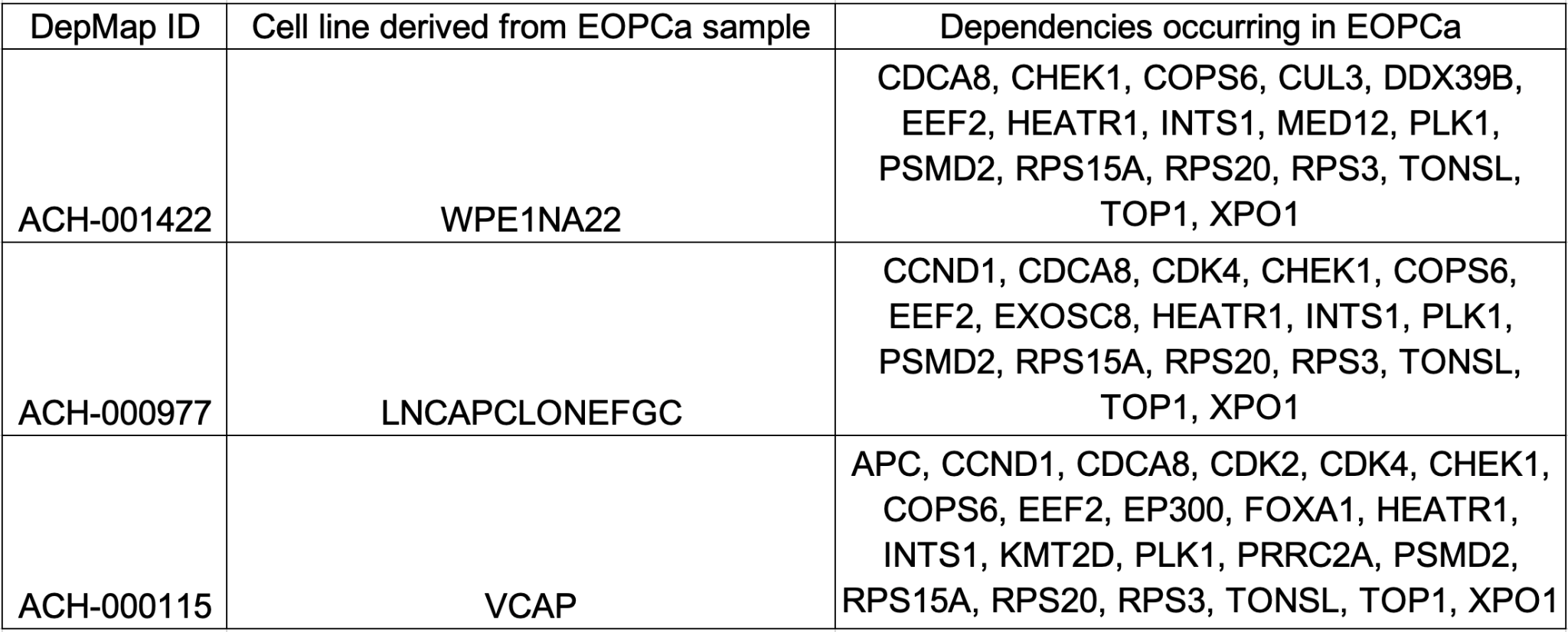

#### Table S4: TCGA Consortium Members

| **First Name** | **Last Name** | **Institution** |
| --- | --- | --- |
| Jaeil | Ahn | Georgetown University Department of Biostatistics, Bioinformatics, and Biomathematics, Washington DC 20057 |
| Rehan | Akbani | The University of Texas MD Anderson Cancer Center, Department of Bioinformatics and Computational Biology, Houston TX 77030 |
| Adrian | Ally | Canada's Michael Smith Genome Sciences Centre, BC Cancer Agency, Vancouver, BC V5Z 4S6, Canada |
| Samirkumar | Amin | SCBMB Graduate Program, Baylor College of Medicine, Houston, TX USA |
| Armen | Aprikian | Urology Dept., MUHC, Montreal General Hospital, 1650 Cedar Avenue, Montreal, QC, H3G 1A4 |
| Joshua | Armenia | Memorial Sloan Kettering Cancer Center, Computational Biology Center, 1275 York Avenue, New York, NY 10065 |
| Arshi | Arora | Department of Epidemiology and Biostatistics, Memorial Sloan-Kettering Cancer Center |
| J. Todd | Auman | Eshelman School of Pharmacy, University of North Carolina at Chapel Hill, Chapel Hill, NC 27599 USA |
| Miruna | Balasundaram | Canada's Michael Smith Genome Sciences Centre, BC Cancer Agency, Vancouver, BC V5Z 4S6, Canada |
| Saianand | Balu | Lineberger Comprehensive Cancer Center, University of North Carolina at Chapel Hill, Chapel Hill, NC 27599 USA |
| Christopher | Benz | 8001 Redwood Blvd., Novato, CA 94945 |
| Alain | Bergeron | Centre de recherche de l'Hôtel-Dieu de Québec, 10, McMahon Street, Québec, QC, G1R 2J6 |
| Rameen | Beroukhim | The Eli and Edythe L. Broad Institute of Massachusetts Institute of Technology and Harvard University Cambridge, Massachusetts 02142, USA. |
| Mario | Berrios | University of Southern California, USC/Norris Comprehensive Cancer Center, 1450 Biggy St., NRT G511, Los Angeles, CA 90033 |
| Tom | Bodenheimer | Lineberger Comprehensive Cancer Center, University of North Carolina at Chapel Hill, Chapel Hill, NC 27599 USA |
| Lori | Boice | UNC Tissue Procurement Facility, Department of Pathology, UNC Lineberger Cancer Center, Chapel Hill, NC 27599, USA |
| Moiz S. | Bootwalla | University of Southern California, USC/Norris Comprehensive Cancer Center, 1450 Biggy St., NRT G511, Los Angeles, CA 90033 |
| Rodolfo | Borges dos Reis | Department of Surgery and Anatomy, Ribeirão Preto Medical School - FMRP, University of São Paulo, Brazil, 14049-900 |
| Paul C. | Boutros | Ontario Institute for Cancer Research, M5G 0A3, Toronto, Ontario, Canada |
| Jay | Bowen | The Research Institute at Nationwide Children's Hospital, Columbus, OH 43205 |
| Reanne | Bowlby | Canada's Michael Smith Genome Sciences Centre, BC Cancer Agency, Vancouver, BC V5Z 4S6, Canada |
| Jeff | Boyd | Fox Chase Cancer Center, 333 Cottman Ave., Philadelphia, PA 19111 |
| Robert K. | Bradley | Computational Biology Program, Public Health Sciences Division and Basic Sciences Division, Fred Hutchinson Cancer Research Center, Seattle WA 98109 |
| Anne | Breggia | Maine Medical Center, 22 Bramhall St, Portland, ME 04102 |
| Fadi | Brimo | Pathology Dept., MUHC, Montreal General Hospital, 1650 Cedar Avenue, Montreal, QC, H3G 1A4 |
| Christopher A. | Bristow | Institute for Applied Cancer Science, Department of Genomic Medicine, The University of Texas MD Anderson Cancer Center, Houston, TX 77030, USA |
| Denise | Brooks | Canada's Michael Smith Genome Sciences Centre, BC Cancer Agency, Vancouver, BC V5Z 4S6, Canada |
| Bradley M. | Broom | The University of Texas MD Anderson Cancer Center, Department of Bioinformatics and Computational Biology, Houston TX 77030 |
| Alan H. | Bryce | Mayo Clinic, 13400 E Shea Blvd, Scottsdale, AZ 85259 |
| Glenn | Bubley | Beth Israel Deaconess medical Center, Boston, MA 02215 |
| Eric | Burks | Lahey Hospital & Medical Center, 41 Mall Road, Burlington, MA 01805 |
| Yaron S.N. | Butterfield | Canada's Michael Smith Genome Sciences Centre, BC Cancer Agency, Vancouver, BC V5Z 4S6, Canada |
| David | Canes | Lahey Hospital & Medical Center, 41 Mall Road, Burlington, MA 01805 |
| Carlos Gilberto | Carlotti Junior | Department of Surgery and Anatomy, Ribeirão Preto Medical School - FMRP, University of São Paulo, Brazil, 14049-900 |
| Rebecca | Carlsen | Canada's Michael Smith Genome Sciences Centre, BC Cancer Agency, Vancouver, BC V5Z 4S6, Canada |
| Michel | Carmel | CHUS, Hôpital Fleurimont, 3001, 12 avenue north, Sherbrooke, QC, J1H 5N4 |
| Peter R. | Carroll | University of CA San Francisco |
| Scott L. | Carter | The Eli and Edythe L. Broad Institute of Massachusetts Institute of Technology and Harvard University Cambridge, Massachusetts 02142, USA. |
| Richard | Cartun | Dept of Pathology, Hartford Hospital, 80 Seymour Street, Hartford, CT 06102 |
| June M. | Chan | University of CA San Francisco |
| Matthew | Chang | Memorial Sloan Kettering Cancer Center, Center for Molecular Oncology, 1275 York Avenue, New York, NY 10065 |
| Yu | Chen | Human Oncology and Pathogenesis Program, and Department of Medicine; Memorial Sloan Kettering Cancer Center, New York, NY 10065 |
| Andrew D. | Cherniack | The Eli and Edythe L. Broad Institute of Massachusetts Institute of Technology and Harvard University Cambridge, Massachusetts 02142, USA. |
| Simone | Chevalier | Research Glen site of the MUHC, Urology Dept., 1001 Decarie Blvd, Montreal, QC, H4A 3J1 |
| Lynda | Chin | Institute for Applied Cancer Science, Department of Genomic Medicine, The University of Texas MD Anderson Cancer Center, Houston, TX 77030, USA |
| Andy | Chu | Canada's Michael Smith Genome Sciences Centre, BC Cancer Agency, Vancouver, BC V5Z 4S6, Canada |
| Eric | Chuah | Canada's Michael Smith Genome Sciences Centre, BC Cancer Agency, Vancouver, BC V5Z 4S6, Canada |
| Sudha | Chudamani | Leidos Biomedical, 9609 Medical Center Dr Rockville, MD 20850 |
| Giovanni | Ciriello | MSKCC, New York, 10065 |
| Amanda | Clarke | Canada's Michael Smith Genome Sciences Centre, BC Cancer Agency, Vancouver, BC V5Z 4S6, Canada |
| Matthew R. | Cooperberg | University of CA San Francisco |
| Niall M. | Corcoran | Department of Surgery, University of Melbourne, Parkville, VIC 3050, Australia |
| Anthony J. | Costello | Epworth Prostate Centre, Richmond, VIC 3121, Australia |
| Janet | Cowan | University of CA San Francisco |
| Daniel | Crain | International Genomics Consortium, 445 N. 5th Street, Phoenix, AZ 85004 |
| Erin | Curley | International Genomics Consortium, 445 N. 5th Street, Phoenix, AZ 85004 |
| Kerstin | David | Indivumed GmbH |
| John A. | Demchok | National Cancer Institute, 31 Center Dr, 3A20, Bethesda, MD 20892 |
| Francesca | Demichelis | Centre for Integrative Biology, University of Trento, Povo Trento 38123, Italy |
| Noreen | Dhalla | Canada's Michael Smith Genome Sciences Centre, BC Cancer Agency, Vancouver, BC V5Z 4S6, Canada |
| Rajiv | Dhir | University of Pittsburgh Pittsburgh, PA 15261 |
| Alexandre | Doueik | Dept. of Pathology, Santa Cabrini Hospital, 6887 Chatelain Street, Montreal, QC, H1T 3X7 |
| Bettina | Drake | Washington University |
| Heidi | Dvinge | Basic Sciences Division, Fred Hutchinson Cancer Research Center, 1100 Fairview Av. N, Seattle, WA 98109 |
| Ina | Felau | National Cancer Institute, 31 Center Dr, 3A20, Bethesda, MD 20892 |
| Martin L. | Ferguson | National Cancer Institute, 31 Center Dr, 3A20, Bethesda, MD 20893 |
| Stephen | Freedland | Cedars Sinai Medical Center, Los Angeles, CA 90048 |
| Yao | Fu | Program in Computational Biology and Bioinformatics, Yale University, Bass 432, 266 Whitney Ave, New Haven, CT 06520 |
| Stacey B. | Gabriel | The Eli and Edythe L. Broad Institute of Massachusetts Institute of Technology and Harvard University Cambridge, Massachusetts 02142, USA. |
| Jianjiong | Gao | Memorial Sloan Kettering Cancer Center, Center for Molecular Oncology, 1275 York Avenue, New York, NY 10065 |
| Johanna | Gardner | International Genomics Consortium, 445 N. 5th Street, Phoenix, AZ 85004 |
| Julie M. | Gastier-Foster | The Research Institute at Nationwide Children's Hospital, Columbus, OH 43205 |
| Mark | Gerken | The Research Institute at Nationwide Children's Hospital, Columbus, OH 43205 |
| Mark | Gerstein | Program in Computational Biology and Bioinformatics, Yale University, Bass 432, 266 Whitney Ave, New Haven, CT 06520 |
| Andrew K. | Godwin | University of Kansas Medical Center, Department of Pathology and Laboratory Medicine, 3901 Rainbow Blvd, Kansas City, KS 66160 |
| Anuradha | Gopalan | Memorial Sloan Kettering Cancer Center, 1275 York Avenue |
| Markus | Graefen | Martini-Clinic, Prostate Cancer Center, University Medical Center Hamburg Eppendorf, Martinistr. 52, 20246 Hamburg, Germany |
| Ranabir | Guin | Canada's Michael Smith Genome Sciences Centre, BC Cancer Agency, Vancouver, BC V5Z 4S6, Canada |
| Manaswi | Gupta | The Eli and Edythe L. Broad Institute of Massachusetts Institute of Technology and Harvard University Cambridge, Massachusetts 02142, USA. |
| Angela | Hadjipanayis | Department of Genetics, Harvard Medical School, Boston, MA 02115, USA |
| Syed | Haider | Ontario Institute for Cancer Research, M5G 0A3, Toronto, Ontario, Canada |
| Lucie | Hamel | Research Glen site of the MUHC, Urology Dept., 1001 Decarie Blvd, Montreal, QC, H4A 3J1 |
| D. Neil | Hayes | Department of Internal Medicine, Division of Medical Oncology, University of North Carolina at Chapel Hill, Chapel Hill, NC 27599 USA |
| David I. | Heiman | The Eli and Edythe L. Broad Institute of Massachusetts Institute Of Technology and Harvard University 415 Main Street Cambridge, MA 02142 |
| Katherine A. | Hoadley | Department of Genetics, University of North Carolina at Chapel Hill, Chapel Hill, NC 27599 USA |
| Andrea H. | Holbrook | University of Southern California, USC/Norris Comprehensive Cancer Center, 1450 Biggy St., NRT G511, Los Angeles, CA 90033 |
| Robert A. | Holt | Canada's Michael Smith Genome Sciences Centre, BC Cancer Agency, Vancouver, BC V5Z 4S6, Canada |
| Antonia | Holway | Lahey Hospital & Medical Center, 41 Mall Road, Burlington, MA 01805 |
| Christopher M. | Hovens | Department of Surgery, University of Melboune, Parkville, VIC 3050 Australia |
| Alan P. | Hoyle | Lineberger Comprehensive Cancer Center, University of North Carolina at Chapel Hill, Chapel Hill, NC 27599 USA |
| Mei | Huang | UNC Tissue Procurement Facility, UNC Lineberger Cancer Center, Chapel Hill, NC 27599, USA |
| Carolyn M. | Hutter | National Human Genome Research Institute, National 20892-9305 Institutes of Health, 5635 Fishers Lane, Bethesda, MD |
| Michael | Ittmann | Baylor College of Medicine |
| Lisa | lype | Institute for Systems Biology, Seattle Washington 98109 |
| Stuart R. | Jefferys | Lineberger Comprehensive Cancer Center, University of North Carolina at Chapel Hill, Chapel Hill, NC 27599 USA |
| Steven J.M. | Jones | Canada's Michael Smith Genome Sciences Centre, BC Cancer Agency, Vancouver, BC V5Z 4S6, Canada |
| Corbin D. | Jones | Department of Biology, University of North Carolina at Chapel Hill, Chapel Hill, NC 27599 USA |
| Hartmut | Juhl | Indivumed GmbH |
| Andre | Kahles | Memorial Sloan Kettering Cancer Center, Computational Biology Center, 1275 York Avenue, New York, NY 10065 |
| Katayoon | Kasaian | Canada's Michael Smith Genome Sciences Centre, BC Cancer Agency, Vancouver, BC V5Z 4S6, Canada |
| Michael | Kerger | Epworth Prostate Centre, Richmond, VIC 3121, Australia |
| Ekta | Khurana | Department of Physiology and Biophysics, Weill Cornell Medical College of Cornell University, 1305 York Avenue, New York, NY 10065 |
| Robert | Klein | Icahn Institute for Genomics and Multiscale Biology, Department of Genetics and Genomic Sciences, Icahn School of Medicine at Mt Sinai, 1 Gustave L. Levy Place, New York, NY 10029 |
| Raju | Kucherlapati | Department of Genetics, Harvard Medical School, Boston, MA 02115, USA |
| Louis | Lacombe | Centre de recherche de l'Hôtel-Dieu de Québec, 10, McMahon Street, Québec, QC, G1R 2J6 |
| Marc | Ladanyi | MSKCC, New York, 10065 |
| Phillip H. | Lai | University of Southern California, USC/Norris Comprehensive Cancer Center, 1450 Biggy St., NRT G511, Los Angeles, CA 90033 |
| Peter W. | Laird | Center for Epigenetics Van Andel Institute, 333 Bostwick Ave N.E. Grand Rapids, MI 49503 |
| Mathieu | Latour | Centre hospitalier St-Luc du CHUM, 1058, rue St-Denis, Montréal, QC, H2X 3J4 |
| Kevin | Lau | International Genomics Consortium, 445 N. 5th Street, Phoenix, AZ 85004 |
| Darlene | Lee | Canada's Michael Smith Genome Sciences Centre, BC Cancer Agency, Vancouver, BC V5Z 4S6, Canada |
| Semin | Lee | Center for Biomedical Informatics, Harvard Medical School, Boston, MA 02115, USA |
| Kjong-Van | Lehmann | Memorial Sloan Kettering Cancer Center, Computational Biology Center, 1275 York Avenue, New York, NY 10065 |
| Kristen M. | Leraas | The Research Institute at Nationwide Children's Hospital, Columbus, OH 43205 |
| Robert | Leung | Weill Cornell Medical College, Department of Urology, New York, NY 10021 |
| John A. | Libertino | Lahey Hospital & Medical Center, 41 Mall Road, Burlington, MA 01805 |
| Tara M. | Lichtenberg | The Research Institute at Nationwide Children's Hospital, Columbus, OH 43205 |
| W. Marston | Linehan | National Cancer Institute, Urologic Oncology Branch, 10 Center Dr, CRC, Rm 1-5940, Bethesda, MD 20892-1107 |
| Shiyun | Ling | The University of Texas MD Anderson Cancer Center, Department of Bioinformatics and Computational Biology, Houston TX 77030 |
| Scott M. | Lippman | UC San Diego Moores Cancer Center, 3855 Health Sciences Drive MC 0685, La Jolla, CA 92093 |
| Wenbin | Liu | The University of Texas MD Anderson Cancer Center, Department of Bioinformatics and Computational Biology, Houston TX 77030 |
| Jia | Liu | Leidos Biomedical, 9609 Medical Center Dr Rockville, MD 20850 |
| Lucas | Lochovsky | Program in Computational Biology and Bioinformatics, Yale University, Bass 432, 266 Whitney Ave, New Haven, CT 06520 |
| Massimo | Loda | Department of Pathology, Dana-Farber Cancer Institute, 450 Brookline Ave, Boston MA 02215 |
| Christopher | Logothetis | The Univeristy of Texas MD Anderson Cancer Center 1515 Holcombe Blvd Unit 1374 Houston. TX 77030 |
| Laxmi | Lolla | Leidos Biomedical, 9609 Medical Center Dr Rockville, MD 20850 |
| Teri | Longacre | Stanford Cancer Institute, Stanford Medicine, 300 Pasteur Drive, Stanford, CA 94305 |
| Yiling | Lu | The University of Texas MD Anderson Cancer Center, Department of Systems Biology, Houston TX 77030 |
| Jianhua | Luo | University of Pittsburgh |
| Yussanne | Ma | Canada's Michael Smith Genome Sciences Centre, BC Cancer Agency, Vancouver, BC V5Z 4S6, Canada |
| Harshad S. | Mahadeshwar | Institute for Applied Cancer Science, Department of Genomic Medicine, The University of Texas MD Anderson Cancer Center, Houston, TX 77030, USA |
| David | Mallery | International Genomics Consortium, 445 N. 5th Street, Phoenix, AZ 85004 |
| Marco A. | Marra | Canada's Michael Smith Genome Sciences Centre, BC Cancer Agency, Vancouver, BC V5Z 4S6, Canada |
| Michael | Mayo | Canada's Michael Smith Genome Sciences Centre, BC Cancer Agency, Vancouver, BC V5Z 4S6, Canada |
| Shannon | McCall | Department of Pathology, Duke Cancer Institute, 200 Trent Drive, Durham, NC 27710 |
| Ginette | McKercher | 1320 Graham Blvd, suite 110, Town of Mount-Royal, QC, H3P 3C8 |
| Shaowu | Meng | Lineberger Comprehensive Cancer Center, University of North Carolina at Chapel Hill, Chapel Hill, NC 27599 USA |
| Maria J. | Merino | Laboratory of Pathology, National Cancer Institute, NIH, Bethesda, MD |
| Anne-Marie | Mes-Masson | CRCHUM, 900 St-Denis Street, Tour Viger, Montreal, QC, H2X 0A9 |
| Matthew | Meyerson | The Eli and Edythe L. Broad Institute of Massachusetts Institute of Technology and Harvard University Cambridge, Massachusetts 02142, USA. |
| Piotr A. | Mieczkowski | Department of Genetics, University of North Carolina at Chapel Hill, Chapel Hill, NC 27599 USA |
| Gordon B. | Mills | The University of Texas MD Anderson Cancer Center, Department of Systems Biology, Houston TX 77030 |
| Sarah | Minner | Dpt. of Pathology, University Medical Center Hamburg Eppendorf, Martinistr. 52, 20246 Hamburg, Germany |
| Alireza | Moinzadeh | Lahey Hospital & Medical Center, 41 Mall Road, Burlington, MA 01805 |
| Richard A. | Moore | Canada's Michael Smith Genome Sciences Centre, BC Cancer Agency, Vancouver, BC V5Z 4S6, Canada |
| Scott | Morris | International Genomics Consortium, 445 N. 5th Street, Phoenix, AZ 85004 |
| Lisle E. | Mose | Lineberger Comprehensive Cancer Center, University of North Carolina at Chapel Hill, Chapel Hill, NC 27599 USA |
| Andrew J. | Mungall | Canada's Michael Smith Genome Sciences Centre, BC Cancer Agency, Vancouver, BC V5Z 4S6, Canada |
| Bradley A. | Murray | The Eli and Edythe L. Broad Institute of Massachusetts Institute of Technology and Harvard University Cambridge, Massachusetts 02142, USA. |
| Jerome B. | Myers | Penrose-St. Francis Health Services Dept. of Pathology, 2222 N. Nevada Ave, Colorado Springs, CO 80907 |
| Rashi | Naresh | SRA International, 4300 Fair Lakes Court Fairfax, VA 22033 |
| Peter S. | Nelson | Division of Human Biology, Fred Hutchinson Cancer Research Center, Seattle, WA 98109 |
| Mark A. | Nelson | University of Arizona |
| Joel | Nelson | University of Pittsburgh Pittsburgh, PA 15261 |
| Houtan | Noushmehr | Department of Genetics, Ribeirão Preto Medical School - FMRP, University of São Paulo, Brazil, 14049-900 |
| Angeliki | Pantazi | Department of Genetics, Harvard Medical School, Boston, MA 02115, USA |
| Michael | Parfenov | Department of Genetics, Harvard Medical School, Boston, MA 02115, USA |
| Peter J. | Park | Center for Biomedical Informatics, Harvard Medical School, Boston, MA 02115, USA |
| Joel S. | Parker | Department of Genetics, University of North Carolina at Chapel Hill, Chapel Hill, NC 27599 USA |
| Joseph | Paulauskis | International Genomics Consortium, 445 N. 5th Street, Phoenix, AZ 85004 |
| Robert | Penny | International Genomics Consortium, 445 N. 5th Street, Phoenix, AZ 85004 |
| Charles M. | Perou | Department of Genetics, University of North Carolina at Chapel Hill, Chapel Hill, NC 27599 USA |
| Alain | Piché | CHUS, Hôpital Fleurimont, 3001, 12 avenue north, Sherbrooke, QC, J1H 5N4 |
| Todd | Pihl | SRA International, 4300 Fair Lakes Court Fairfax, VA 22033 |
| Peter A. | Pinto | National Cancer Institute, Urologic Oncology Branch, 10 Center Dr, CRC, Rm 1-5940, Bethesda, MD 20892-1107 |
| Davide | Prandi | Centre for Integrative Biology, University of Trento, Povo Trento 38123, Italy |
| Alexei | Protopopov | Institute for Applied Cancer Science, Department of Genomic Medicine, The University of Texas MD Anderson Cancer Center, Houston, TX 77030, USA |
| Kenna | R. Mills Shaw | MD Anderson Cancer Center, Houston, TX 77230 |
| Nilsa C. | Ramirez | The Research Institute at Nationwide Children's Hospital, Columbus, OH 43205 |
| Arvind | Rao | The University of Texas MD Anderson Cancer Center, Department of Bioinformatics and Computational Biology, Houston TX 77030 |
| W. K. | Rathmell | Lineberger Comprehensive Cancer Center, University of North Carolina at Chapel Hill, Chapel Hill, NC 27599 USA |
| Gunnar | Rätsch | Memorial Sloan Kettering Cancer Center, Computational Biology Center, 1275 York Avenue, New York, NY 10065 |
| Xiaojia | Ren | Department of Genetics, Harvard Medical School, Boston, MA 02115, USA |
| Victor | Reuter | Memorial Sloan Kettering Cancer Center, Department of Pathology, 1275 York Avenue, New York, NY 10065 |
| Sheila M. | Reynolds | Institute for Systems Biology, Seattle Washington 98109 |
| Suhn K. | Rhie | Norris Comprehensive Cancer Center Keck School of Medicine, University of Southern California, 1441 Eastlake Avenue, Los Angeles, CA 90033 |
| Kimberly | Rieger-Christ | Lahey Hospital & Medical Center, 41 Mall Road, Burlington, MA 01805 |
| Jeffrey | Roach | Research Computing Center, University of North Carolina at Chapel Hill, Chapel Hill, NC 27599 USA |
| A. Gordon | Robertson | Canada's Michael Smith Genome Sciences Centre, BC Cancer Agency, Vancouver, BC V5Z 4S6, Canada |
| Brian | Robinson | Weill Cornell Medical College, Department of Pathology & Laboratory Medicine, 1300 York Avenue, New York, NY 10065 |
| Mark A. | Rubin | Weill Cornell Medicial College of Cornell University |
| Fred | Saad | CRCHUM, 900 St-Denis Street, Tour Viger, Montreal, QC, H2X 0A9 |
| Sara | Sadeghi | Canada's Michael Smith Genome Sciences Centre, BC Cancer Agency, Vancouver, BC V5Z 4S6, Canada |
| Gordon | Saksena | The Eli and Edythe L. Broad Institute of Massachusetts Institute of Technology and Harvard University Cambridge, Massachusetts 02142, USA. |
| Charles | Saller | Analytical Biological Services, 701 Cornell Drive, Wilmington, DE 19801 |
| Andrew | Salner | Gray Cancer Center, Hartford Hospital, 80 Seymour Street, Hartford, CT 06102 |
| Chris | Sander | Dana-Farber Cancer Institute, 450 Brookline Ave, Boston MA 02215 |
| George | Sandusky | Indiana University, Dept Pathology, Med A Science Bldg Rm 128, 635 Barnhill Drive, Indianpolis |
| Guido | Sauter | Dpt. of Pathology, University Medical Center Hamburg Eppendorf, Martinistr. 52, 20246 Hamburg, Germany |
| Andrea | Sboner | Weill Cornell Medicial College of Cornell University; Department of Pathology and Laboratory Medicine, Institute for Computational Biomedicine, Institute for Precision Medicine, 1305 York Ave, New York, NY 10021 |
| Eleonora | Scarlata | Research Glen site of the MUHC, Urology Dept., 1001 Decarie Blvd, Montreal, QC, H4A 3J1 |
| Jacqueline E. | Schein | Canada's Michael Smith Genome Sciences Centre, BC Cancer Agency, Vancouver, BC V5Z 4S6, Canada |
| Thorsten | Schlomm | Martini-Clinic, Prostate Cancer Center, University Medical Center Hamburg Eppendorf, Martinistr. 52, 20246 Hamburg, Germany |
| Laura S. | Schmidt | Urologic Oncology Branch, Center for Cancer Research, National Cancer Institute, NIH, Bethesda MD and Basic Science Program, Leidos Biomedical Research, Inc, Frederick National Laboratory for Cancer Research, Frederick, MD |
| Nikolaus | Schultz | Memorial Sloan Kettering Cancer Center, Center for Molecular Oncology, 1275 York Avenue, New York, NY 10065 |
| Steven E. | Schumacher | The Eli and Edythe L. Broad Institute of Massachusetts Institute of Technology and Harvard University Cambridge, Massachusetts 02142, USA. |
| Jonathan | Seidman | Department of Genetics, Harvard Medical School, Boston, MA 02115, USA |
| Luciano Neder | Serafini | Department of Pathology, Ribeirão Preto Medical School - FMRP, University of São Paulo, Brazil, 14049-900 |
| Sahil | Seth | Institute for Applied Cancer Science, Department of Genomic Medicine, The University of Texas MD Anderson Cancer Center, Houston, TX 77030, USA |
| Alexis | Sharp | Department of Pathology, Duke Cancer Institute, 200 Trent Drive, Durham, NC 27710 |
| Troy | Shelton | International Genomics Consortium, 445 N. 5th Street, Phoenix, AZ 85004 |
| Candace | Shelton | International Genomics Consortium, 445 N. 5th Street, Phoenix, AZ 85004 |
| Hui | Shen | Center for Epigenetics, Van Andel Institute, 333 Bostwick Ave N.E. Grand Rapids, MI 149503 |
| Ronglai | Shen | Department of Epidemiology and Biostatistics, Memorial Sloan-Kettering Cancer Center |
| Mark | Sherman | International Genomics Consortium, 445 N. 5th Street, Phoenix, AZ 85004 |
| Margi | Sheth | National Cancer Institute, 31 Center Dr, 3A20, Bethesda, MD 20892 |
| Yan | Shi | Lineberger Comprehensive Cancer Center, University of North Carolina at Chapel Hill, Chapel Hill, NC 27599 USA |
| Juliann | Shih | The Eli and Edythe L. Broad Institute of Massachusetts Institute of Technology and Harvard University Cambridge, Massachusetts 02142, USA. |
| Ilya | Shmulevich | Institute for Systems Biology, Seattle Washington 98109 |
| Jeffry | Simko | University of CA San Francisco |
| Ronald | Simon | Dpt. of Pathology, University Medical Center Hamburg Eppendorf, Martinistr. 52, 20246 Hamburg, Germany |
| Janae V. | Simons | Lineberger Comprehensive Cancer Center, University of North Carolina at Chapel Hill, Chapel Hill, NC 27599 USA |
| Payal | Sipahimalani | Canada's Michael Smith Genome Sciences Centre, BC Cancer Agency, Vancouver, BC V5Z 4S6, Canada |
| Tara | Skelly | Department of Genetics, University of North Carolina at Chapel Hill, Chapel Hill, NC 27599 USA |
| Heidi J. | Sofia | National Human Genome Research Institute, National Institutes of Health, 5635 Fishers Lane, Bethesda, MD 20892-9305 |
| Matthew G. | Soloway | Lineberger Comprehensive Cancer Center, University of North Carolina at Chapel Hill, Chapel Hill, NC 27599 USA |
| Xingzhi | Song | Institute for Applied Cancer Science, Department of Genomic Medicine, The University of Texas MD Anderson Cancer Center, Houston, TX 77030, USA |
| Andrea | Sorcini | Lahey Hospital & Medical Center, 41 Mall Road, Burlington, MA 01805 |
| Carrie | Sougnez | The Eli and Edythe L. Broad Institute of Massachusetts Institute of Technology and Harvard University Cambridge, Massachusetts 02142, USA. |
| John | Stewart | Wake Forest Baptist Health, 1 Medical Center Boulevard, Winston-Salem, North Carolina, 27157 |
| Travis B. | Sullivan | Lahey Hospital & Medical Center, 41 Mall Road, Burlington, MA 01805 |
| Huandong | Sun | Institute for Applied Cancer Science, Department of Genomic Medicine, The University of Texas MD Anderson Cancer Center, Houston, TX 77030, USA |
| Charlie | Sun | SRA International, 4300 Fair Lakes Court Fairfax, VA 22033 |
| Angela | Tam | Canada's Michael Smith Genome Sciences Centre, BC Cancer Agency, Vancouver, BC V5Z 4S6, Canada |
| Donghui | Tan | Department of Genetics, University of North Carolina at Chapel Hill, Chapel Hill, NC 27599 USA |
| Jiabin | Tang | Institute for Applied Cancer Science, Department of Genomic Medicine, The University of Texas MD Anderson Cancer Center, Houston, TX 77030, USA |
| Roy | Tarnuzzer | National Cancer Institute, 31 Center Dr, 3A20, Bethesda, MD 20892 |
| Katherine | Tarvin | Analytical Biological Services 701 Cornell Drive Wilmington, DE 19801 |
| Barry S. | Taylor | Human Oncology and Pathogenesis Program, Department of Epidemiology and Biostatistics, and Center for Molecular Oncology; Memorial Sloan Kettering Cancer Center, New York, NY 10065 |
| Patrick | Teebagy | Lahey Hospital & Medical Center, 41 Mall Road, Burlington, MA 01805 |
| Imelda | Tenggara | University of CA San Francisco |
| Bernard | Têtu | Hôpital du St-Sacrement, CHU de Québec, 1050, chemin Ste-Foy, Québec QC, G1S 4L8 |
| Ashutosh | Tewari | Weill Cornell Medical College, Department of Urology, New York, NY 10021 |
| Nina | Thiessen | Canada's Michael Smith Genome Sciences Centre, BC Cancer Agency, Vancouver, BC V5Z 4S6, Canada |
| Timothy | Thompson | The Univeristy of Texas MD Anderson Cancer Center; 1515 Holcombe Blvd Unit 0018-3, Houston, TX 77030 |
| Leigh B. | Thorne | UNC Tissue Procurement Facility, Department of Pathology, UNC Lineberger Cancer Center, Chapel Hill, NC 27599, USA |
| Daniela Pretti da Cunha | Tirapelli | Department of Surgery and Anatomy, Ribeirão Preto Medical School - FMRP, University of São Paulo, Brazil, 14049-900 |
| Scott A. | Tomlins | University of Michigan Medical School |
| Felipe Amstalden | Trevisan | Department of Surgery and Anatomy, Ribeirão Preto Medical School - FMRP, University of São Paulo, Brazil, 14049-900 |
| Patricia | Troncoso | The Univeristy of Texas MD Anderson Cancer Center 1515 Holcombe Blvd Unit 0085 Houston, TX 77030 |
| Lawrence | True | University of Washington, 1959 NE Pacific St., Seattle, WA, 98195-6100 |
| Maria Christina | Tsourlakis | Dpt. of Pathology, University Medical Center Hamburg Eppendorf, Martinistr. 52, 20246 Hamburg, Germany |
| Svitlana | Tyekucheva | Dana-Farber Cancer Institute, 450 Brookline Ave, Boston MA 02215 |
| Eliezer | Van Allen | Broad Institute of MIT and Harvard 415 Main Street Cambridge, MA 02142 |
| David J. | Van Den Berg | University of Southern California, USC/Norris Comprehensive Cancer Center, 1450 Biggy St., NRT G511, Los Angeles, CA 90033 |
| Umadevi | Veluvolu | Department of Genetics, University of North Carolina at Chapel Hill, Chapel Hill, NC 27599 USA |
| Roel | Verhaak | The University of Texas MD Anderson Cancer Center, Department of Bioinformatics and Computational Biology, Houston TX 77030 |
| Cathy D. | Vocke | Urologic Oncology Branch, Center for Cancer Research, National Cancer Institute, NIH, Bethesda MD |
| Yunhu | Wan | SRA International, 4300 Fair Lakes Court, Fairfax, VA 22033 |
| Zhining | Wang | National Cancer Institute, 31 Center Dr, 3A20, Bethesda, MD 20892 |
| Qingguo | Wang | Memorial Sloan Kettering Cancer Center, Center for Molecular Oncology, 1275 York Avenue, New York, NY 10065 |
| Wenyi | Wang | The University of Texas MD Anderson Cancer Center, Department of Bioinformatics and Computational Biology, Houston, TX 77030 |
| Nils | Weinhold | Memorial Sloan Kettering Cancer Center, Computational Biology Center, 1275 York Avenue, New York, NY 10065 |
| John N. | Weinstein | The University of Texas MD Anderson Cancer Center, Department of Bioinformatics and Computational Biology, Houston TX 77030 |
| Daniel J. | Weisenberger | University of Southern California, USC/Norris Comprehensive Cancer Center, 1450 Biggy St., NRT G511, Los Angeles, CA 90033 |
| Matthew D. | Wilkerson | Department of Genetics, University of North Carolina at Chapel Hill, Chapel Hill, NC 27599 USA |
| Lisa | Wise | The Research Institute at Nationwide Children's Hospital, Columbus, OH 43205 |
| John | Witte | University of CA San Francisco |
| Chia-Chin | Wu | Department of Genomic Medicine, The University of Texas MD Anderson Cancer Center. Houston, TX 77030 USA |
| Junyuan | Wu | Lineberger Comprehensive Cancer Center, University of North Carolina at Chapel Hill, Chapel Hill, NC 27599 USA |
| Ye | Wu | Leidos Biomedical, 9609 Medical Center Dr Rockville, MD 20850 |
| Andrew W. | Xu | Center for Biomedical Informatics, Harvard Medical School, Boston, MA 02115, USA |
| Shalini S. | Yadav | Weill Cornell Medical College, Department of Urology, New York, NY 10021 |
| Lixing | Yang | Center for Biomedical Informatics, Harvard Medical School, Boston, MA 02115, USA |
| Liming | Yang | National Cancer Institute, 31 Center Dr, 3A20, Bethesda, MD 20892 |
| Christina | Yau | Buck Institute |
| Huihui | Ye | Beth Israel Deaconess Medical Center, 330 Brookline Ave, Boston MA 02215 |
| Peggy | Yena | International Genomics Consortium, 445 N. 5th Street, Phoenix, AZ 85004 |
| Thomas | Zeng | Canada's Michael Smith Genome Sciences Centre, BC Cancer Agency, Vancouver, BC V5Z 4S6, Canada |
| Jean C. | Zenklusen | National Cancer Institute, 31 Center Dr, 3A20, Bethesda, MD 20892 |
| Jianhua | Zhang | Institute for Applied Cancer Science, Department of Genomic Medicine, The University of Texas MD Anderson Cancer Center, Houston, TX 77030, USA |
| Wei | Zhang | Department of Pathology, University of Texas MD Anderson Cancer Center, Texas 77030 |
| Jiashan | Zhang | National Cancer Institute, 31 Center Dr, 3A20, Bethesda, MD 20892 |
| Yi | Zhong | Memorial Sloan Kettering Cancer Center, Computational Biology Center, 1275 York Avenue, New York, NY 10065 |
| Kelsey | Zhu | Canada's Michael Smith Genome Sciences Centre, BC Cancer Agency, Vancouver, BC V5Z 4S6, Canada |
| Erik | Zmuda | The Research Institute at Nationwide Children's Hospital, Columbus, OH 43205 |

#### Table S5: ICGC Consortium Members

| **First Name** | **Last Name** | **Institution** |
| --- | --- | --- |
| Abraham | Gihawi | Norwich Medical School, University of East Anglia, Norwich, NR4 7TJ, UK |
| Adam | Butler | The Cancer, Ageing and Somatic Mutation Programme, Wellcome Trust Sanger Institute, Hinxton, Cambridge, CB10 1SA, UK |
| Adam | Lambert | The University of Oxford, Oxford, OX1 2JD, UK |
| Andy | Lynch | School of Mathematics and Statistics/School of Medicine, University of St Andrews, St Andrews, Fife, KY16 9SS, UK; Cancer Research UK Cambridge Institute, University of Cambridge, Li Ka Shing Centre, Robinson Way, Cambridge, CB2 0RE, UK |
| Anne | Warren | Cambridge University Hospitals NHS FT & University of Cambridge, Cambridge Biomedical Campus, Cambridge, CB2 0QQ, UK |
| Atef | Sahil | Big Data Institute, University of Oxford, Old Road Campus, Oxford, OX3 7LF, UK |
| Charlie | Massie | Department of Oncology, Hutchison/MRC Research Centre, Cambridge University, Cambridge, CB2 0XZ, UK; Early Detection Programme, CRUK Cambridge Centre, Cancer Research UK Cambridge Institute, Robinson Way, Cambridge, CB2 ORE, UK |
| Christopher | Foster | The Institute of Cancer Research, London, SW7 3RP, UK |
| Clare | Verrill | Nuffield Department of Surgical Sciences, University of Oxford, Headington, Oxford, OX3 9DU, UK; Oxford NIHR Biomedical Research Centre, Oxford, UK |
| Colin | Cooper | The Institute of Cancer Research, London, SW7 3RP, UK |
| Dan | Berney | Department of Molecular Oncology, Barts Cancer Centre, Barts and the London School of Medicine and Dentistry, London, E1 2AD, UK |
| Dan | Burns | The Institute of Cancer Research, London, SW7 3RP, UK |
| Dan | Woodcock | Big Data Institute, University of Oxford, Old Road Campus, Oxford, OX3 7LF, UK |
| Daniel | Brewer | Norwich Medical School, University of East Anglia, Norwich, NR4 7TJ, UK |
| David | Wedge | The University of Manchester, Manchester, M13 9PL, UK; Big Data Institute, University of Oxford, Old Road Campus, Oxford, OX3 7LF, UK |
| Elizabeth | Bancroft | The Institute of Cancer Research, London, SW7 3RP, UK; Royal Marsden NHS Foundation Trust, London and Sutton, SM2 5PT, UK |
| Freddie | Hamdy | The University of Oxford, Oxford, OX1 2JD, UK |
| Steven | Bova | Institute of Biosciences and Medical Technology, BioMediTech, University of Tampere and Fimlab Laboratories, Tampere University Hospital, Tampere, FI-33520, Finland |
| Gahee | Park | Department of Oncology, Hutchison/MRC Research Centre, Cambridge University, Cambridge, CB2 0XZ, UK |
| Gregory | Leeman | The Cancer, Ageing and Somatic Mutation Programme, Wellcome Trust Sanger Institute, Hinxton, Cambridge, CB10 1SA, UK |
| Harveer | Dev | Early Cancer Institute, University of Cambridge, Department of Oncology, Box 197, Cambridge Biomedical Campus, Cambridge, CB2 0XZ, UK |
| Ian | Mills | Nuffield Department of Surgical Sciences, University of Oxford, UK |
| Jingjing | Zhang | Norwich Medical School, University of East Anglia, Norwich, NR4 7TJ, UK |
| Melissa | Cheung | Early Cancer Institute, University of Cambridge, Department of Oncology, Box 197, Cambridge Biomedical Campus, Cambridge, CB2 0XZ, UK |
| Nening | Dennis | Royal Marsden NHS Foundation Trust, London and Sutton, SM2 5PT, UK |
| Peter | Campbell | The Cancer, Ageing and Somatic Mutation Programme, Wellcome Trust Sanger Institute, Hinxton, Cambridge, CB10 1SA, UK |
| Peter | Van Loo | The Francis Crick Institute, Department of Human Genetics, London, NW1 1AT, UK; The University of Texas, M.D. Anderson Cancer Center, 1515 Holcombe Boulevard, Houston, Texas, 77030, USA |
| Radoslaw | Lach | Department of Oncology, Hutchison/MRC Research Centre, Cambridge University, Cambridge, CB2 0XZ, UK |
| Rosalind | Eeles | The Institute of Cancer Research, London, SW7 3RP, UK; Royal Marsden NHS Foundation Trust, London and Sutton, SM2 5PT, UK |
| Sara | Pita | Department of Oncology, Hutchison/MRC Research Centre, Cambridge University, Cambridge, CB2 0XZ, UK |
| Steven | Hazell | Royal Marsden NHS Foundation Trust, London and Sutton, SM2 5PT, UK |
| Sue | Merson | The Institute of Cancer Research, London, SW7 3RP, UK |
| Tapio | Visakorpi | Finland bioinformatics, Institute of Biosciences and Medical Technology, BioMediTech, University of Tampere and Fimlab Laboratories, Tampere University Hospital, Tampere, FI-33520, Finland |
| Thomas | Mitchell | The Cancer, Ageing and Somatic Mutation Programme, Wellcome Trust Sanger Institute, Hinxton, Cambridge, CB10 1SA, UK; Department of Urology, Addenbrooke’s Hospital, Cambridge, CB2 0QQ, UK; Academic Urology Group, University of Cambridge, Cambridge, CB2 0RE, UK |
| Toby | Milne-Clark | Early Cancer Institute, University of Cambridge, Department of Oncology, Box 197, Cambridge Biomedical Campus, Cambridge, CB2 0XZ, UK |
| Tokhir | Dadaev | The Institute of Cancer Research, London, SW7 3RP, UK |
| Valeriia | Haberland | Norwich Medical School, University of East Anglia, Norwich, NR4 7TJ, UK |
| Vincent | Gnanapragasam | Department of Urology, Addenbrooke’s Hospital, Cambridge, CB2 0QQ, UK; Academic Urology Group, University of Cambridge, Cambridge, CB2 0RE, UK |
| Yaobo | Xu | The Cancer, Ageing and Somatic Mutation Programme, Wellcome Trust Sanger Institute, Hinxton, Cambridge, CB10 1SA, UK |
| Yong | Jie-Lu | Department of Molecular Oncology, Barts Cancer Centre, Queen Mary University of London, London, EC1M 6BD, UK; First Affiliated Hospital, Zhengzhou University, Zhengzhou, China |
| Zsofia | Kote-Jarai | The Institute of Cancer Research, London, SW7 3RP, UK |
